# The anxiolytic-like effect of the aqueous extract of *Punica granatum* depends on the stress level of fish and mice: Evidence of a homeostatic re-establishing effect

**DOI:** 10.64898/2026.10.08.757595

**Authors:** Aurora Rúbria Batista Pantoja, Jessica Montserrat Chan-Monroy, Marissol Leite, Nancy Cervantes-Anaya, Nelly Maritza Vega-Rivera, Thaeny Regina Ribeiro Rodrigues, Adriano Oliveira Silva Meneses, Bruna Patrícia Dutra Costa, Diógenes Henrique de Siqueira-Silva, Monica Lima-Maximino, Erika Estrada-Camarena, Caio Maximino

## Abstract

Pomegranate (*Punica granatum*) has shown potential anxiolytic and antidepressant effects in preclinical research, but its mechanisms remain unclear. Importantly, some evidence suggests that the effects of *P. granatum* depend on previous stress, suggesting an adaptogenic effect. This study aimed to evaluate the anxiolytic-like effects of an aqueous extract of *P. granatum* (AE-PG) on lambaris and zebrafish exposed to acute restraint stress (ARS) and female BALB/c mice exposed to unpredictable chronic mild stress (CUMS). In addition, we evaluated antidepressant-like effects in female BALB/c mice. Acute administration of AE-PG (1 mg/kg, i.p.) was tested in the novel tank test (NTT) in lambaris and zebrafish and in the light/dark test (LDT) in zebrafish, while chronic (7 days) administration of AE-PG (1 mg/kg, i.p.) was tested in the elevated plus-maze (EPM), open-field test (OFT), splash test (ST), and forced swim test (FST) in mice. ARS increased geotaxis and erratic swimming in the NTT, as well as scototaxis, risk assessment, and erratic swimming in the LDT. Acute pre-treatment with AE-PG blocked the effects of ARS on these anxiety-like endpoints in both lambaris and zebrafish. CUMS increased open arm avoidance in the EPM, thigmotaxis and horizontal and vertical exploration in the OFT, and immobility in the FST, and decreased grooming in the ST. Chronic treatment with AE-PG blocked the effects of CUMS on anxiety-like endpoints, and depression-like endpoints, in mice. An *in silico* analysis of the main compounds in AE-PG suggested a possible mechanism of action on a wide network of targets that converge on PI3K/Akt and show differential expression in brain regions associated with fear- and anxiety-like behavior. Molecular docking suggests that a compound from AE-PG, ellagic acid, is likely to act through estrogen receptors, which could help reestablish homeostatic balance in response to stress.

## 1. Introduction

Among psychiatric disorders, those with the highest prevalence fall in the groups of anxiety, mood, and common mental health disorders (which include nonspecific symptoms of anxiety, depression, and somatic complaints) (J. Liu et al., 2024; Q. Liu et al., 2020; Steel et al., 2014; Yang et al., 2021). Nonetheless, although current pharmacological treatments reduce symptoms, there is still room for improvement, as effects are usually small (Huhn et al., 2014; Kopcalic et al., 2025). The use of medicinal plants as treatments or coadjuvants in the treatment of these disorders emerges as an important area of inquiry, intersecting between the fields of behavioral neuroscience, ethnopharmacology, and medicinal chemistry (Kaushal et al., 2024).

Among the plants studied in preclinical settings is pomegranate (*Punica granatum*), a fruit-bearing deciduous shrub in the family Lythracea that was first domesticated in the region spanning the Caucasus and the Iranian plateau (C. Liu et al., 2025). References to sacred, mythological, and medicinal uses of pomegranate abound in the cultures to which it spread (Alp, 2025). Pomegranate (PG) extracts contain dozens of secondary metabolites across peel, seed, flowers, and leaves, and no single compound has been definitively established as responsible for the observed pharmacological effects (C. Liu et al., 2025). The include ellagitannins, especially punicalagin and punicalin, which are major polyphenols concentrated in the peel and juice; ellagic acid, produced as ellagitannins break down, and present in the arils, juice, and peel; anthocyanins, including cyanidin, delphinidin, and pelargonidin glycosides; other flavonoids, including derivatives of quercetin, kaempferol, and luteolin; phenolic acids such as gallic acid, caffeic acid, and chlorogenic acid; other tannins, including gallotannins and related hydrolysable tannins, particularly abundant in the peel; alkaloids, including pelletierine and isopelletierine, found mainly in the peel and root bark; triterpenes and sterols, including ursolic acid and oleanolic acid, chiefly in the peel and seed oil; and seed-oil fatty acids (notably punicic acid, a conjugated linolenic acid)(C. Liu et al., 2025). Some of these compounds – notably ellagitannins and ellagic acid – have been implicated in the neurobehavioral effects of pomegranate extracts in animal models (Cervantes-Anaya et al., 2022; Estrada-Camarena et al., 2020; González-Trujano et al., 2015).

PG extracts have been patented to be used as an adjuvant in treating anxiety and depression disorders, especially in menopausal women (H. R. Lee, 2010/2012; H.-Y. Lee, 2010/2011). While it shows preclinical promise for reducing anxiety- and depression-like behaviors, no clinical trials testing these effects in humans exist, and nearly all evidence derives from rodent or zebrafish models (Flores-Bazán et al., 2023). In rodents, an aqueous extract of whole-fruit PG (AE-PG) has been shown to produce antinociceptive and anti-inflammatory effects (González-Trujano et al., 2015), antidepressant- and anxiolytic-like effects in ovariectomized (OVX) female rats (Cervantes-Anaya et al., 2022; Estrada-Camarena et al., 2020; Valdés-Sustaita et al., 2017, 2021), and to reduce ischemia-induced anxiety-like behavior in male rats (Rafieirad & Abbaszadeh, 2017). Ethanolic extracts of PG peel (200 mg/kg) appear to produce antidepressant-like effects in male rats, with efficacy comparable to imipramine (Syeda et al., 2020). In a chronic fatigue model in which male mice are subjected to forced swimming for 6 days, the ethanolic leaf extract (100 and 200 mg/kg) reduces immobility (Devi & Das, 2016). AE-PG appears to produce antidepressant-like effects at much smaller doses (1.0-10.0 mg/kg) in OVX rats (Cervantes-Anaya et al 2022; Valdés-Sustaita et al., 2017; Vega-Rivera et al., 2023). Chronic co-treatment with AE-PG (0.125 mg/kg) facilitates the antidepressant response of citalopram at sub-threshold doses in OVX rats, increasing dendritic complexity, dendritic spine number, and expression of mature BDNF and synapthophysin in the dentate gyrus (Vega-Rivera et al., 2023).

The anxiolytic-like effects of PG are less well-studied; PG juice at 20% and 40% concentrations inhibited aluminum chloride-induced anxiogenic effects across elevated plus maze, light/dark box, and open-field tests (Abu-Taweel & Al-Mutary, 2021). Chronic treatment with ethanolic PG seed extract (400 mg/kg) significantly improved ischemia-induced anxiety and locomotor deficits in rats subjected to bilateral carotid ligation (Rafieirad & Abbaszadeh, 2017). Ethanolic leaf extract reversed stress-induced anxiety in the elevated plus maze and mirror chamber tests, an effect attributed to flavonoid modulation of GABA_A_ receptors (Devi & Das, 2016). Chronic treatment with AE-PG (2 mg/kg) prevented anxiety-like behavior induced by a cafeteria diet in OVX rats, associated with modulation of peroxisome proliferator-activated receptor γ and extracellular signal-regulated protein kinase 1/2 activity in the amygdala and hippocampus (Estrada-Camarena et al., 2020).

The fact that AE-PG produces clear effects in OVX rats suggests mediation by estrogen receptors (Estrada-Camarena et al., 2024; Valdés-Sustaita et al., 2017, 2021). Other mechanisms have also been proposed, including the serotonin and kynurenine pathways (Jahromy et al., 2014; Zhu et al., 2021), nitric oxide signaling (Cervantes-Anaya et al., 2025), and the TLR4/NF-kB anti-inflammatory pathway (P. Chen et al., 2024). Ellagic acid and punicalagin, two compounds found in AE-PG, appear to act through different monoaminergic systems (Aleksandrova et al., 2023); ellagic acid preferentially affects swimming (serotonergic) behavior while punicalagin influences climbing (noradrenergic) behavior (Abdel Moneim, 2012; Cervantes-Anaya et al., 2022; Dhingra & Chhillar, 2012; Loren et al., 2005), yet the discrepancy may stem from varying polyphenol ratios in different extracts. Quercetin, an antioxidant flavonoid found in pomegranate, may produce an anxiolytic-like effect in Swiss mice (Islam et al., 2022). This is achieved through binding to GABA_A_ α_5_ subunits, based on results obtained from molecular docking and behavioral tests (Islam et al., 2022)(Islam et al., 2022).

The many putative targets of pomegranate-derived molecules suggest that this plant can be used as an “adaptogen”, a stress-response modifiers that increase an organism’s nonspecific resistance to stress (Panossian, 2017). Adaptogens exhibit multitarget action, which includes receptors for corticosteroid, mineralocorticoid, progestin, estrogen, serotonin (5-HT), N-methyl-*D*-aspartate, and acetylcholine (Panossian, 2017). Interestingly, prior stress exposure is a predominant experimental context in preclinical research with PG, as most studies induced behavioral deficits with a specific stressor before testing extract efficacy as an antidepressant or anxiolytic (Estrada-Camarena et al., 2024). In naive mice without prior stress, peel extract alone produced antidepressant effects in forced swim test (FST) (Syeda et al., 2020), whereas in non-stressed rats, the effect in anxiety is controversial and highly dose-dependent (Estrada-Camarena et al 2020; 2024). The effects in the FST are consistent with the need for prior stress, as swimming is a strong stressor in rodents, revealing a behavioral effect even in naive animals. Oxidative stress is a convergent mechanistic target, with pomegranate extracts reducing lipid peroxidation and enhancing antioxidant enzymes (catalase, SOD, GST, GSH) across multiple models (Abu-Taweel & Al-Mutary, 2021; Cervantes-Anaya et al., 2022, 2025; Devi & Das, 2016). These findings suggest pomegranate extracts ameliorate stress-induced behavioral deficits primarily through antioxidant and neurotransmitter-restorative mechanisms, though direct comparisons across extract types and stress paradigms remain limited.

In the present work, we evaluated the role of previous stress (acute and chronic) in the behavioral effects of AE-PG (1 mg/kg) in lambaris (*Astyanax bimaculatus*), zebrafish (*Danio rerio*), and mice (*Mus musculus*). We found that both acute and chronic stress induced an anxiogenic-like profile in all three species, and those effects were blocked by AE-PG; however, the effects of AE-PG in non-stressed animals are lost. The antidepressant-like effect remains independent of previous stress exposure. Using bioinformatic approaches, we show that AE-PG acts at a wide network of targets that converge on PI3K/Akt and are differentially expressed in brain regions associated with fear- and anxiety-like behavior, and that a compound from AE-PG, ellagic acid, likely acts through estrogen receptors.

## 2. Methods

### 2.1. Aqueous extract preparation

The AE-PG was donated by Nutraceuticals S. L.(Elche, Alicante Spain) prepared from the whole fruit and dissolved in saline solution (0.9 %). The dose selection was based on a previous study in which AE-PG showed an antidepressant and anxiolytic-like effect in ovariectomized rats (Estrada-Camarena et al., 2020; Valdés-Sustaita et al., 2017). Table 1 shows the concentration of the main polyphenolic compounds in AE-PG, as previously reported by Gonzalez-Trujano et al. (2015) and confirmed by Cervantes-Anaya et al. (2022).

**Table 1.**
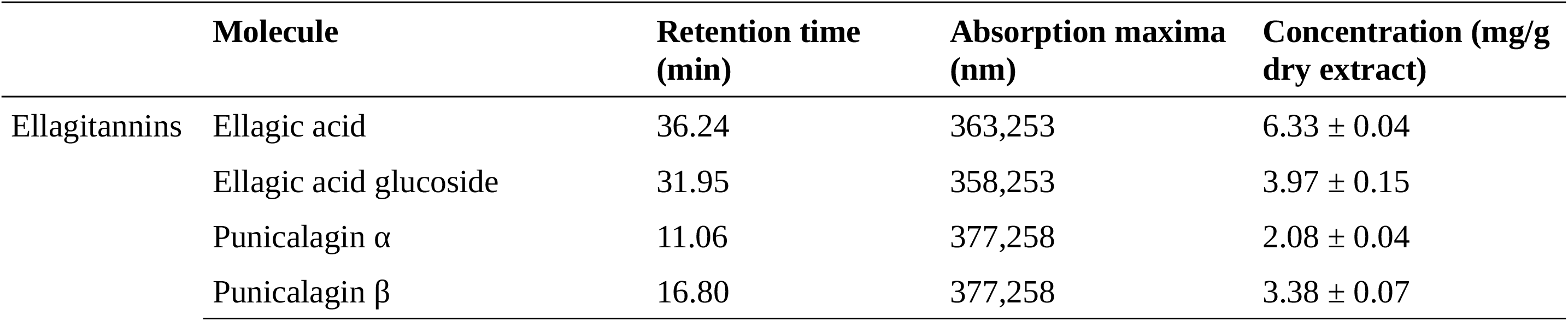
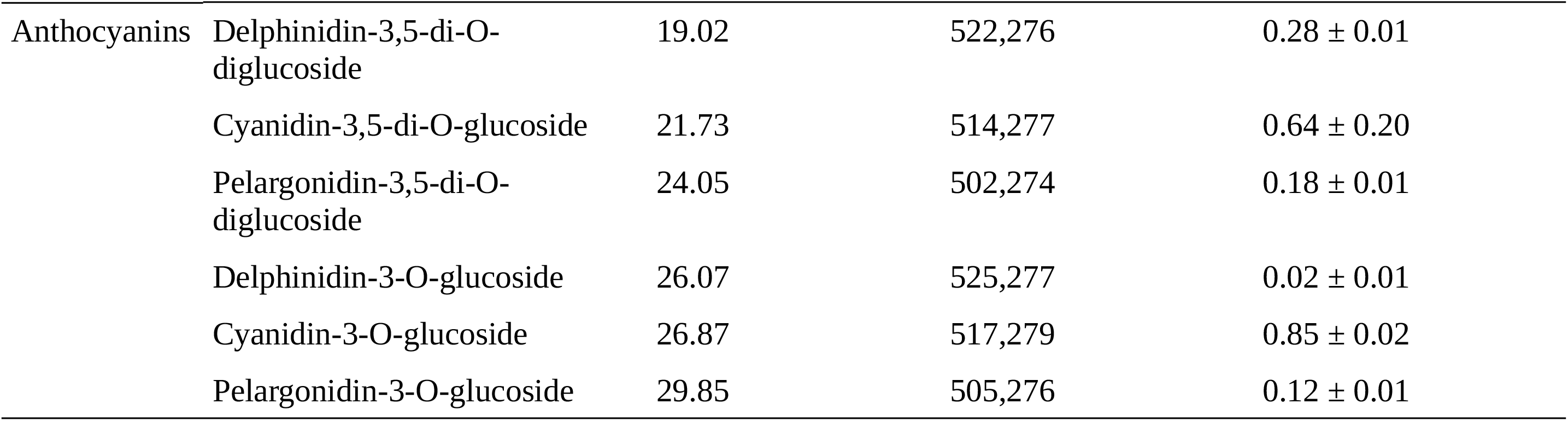
Concentration of main polyphenolic compounds in the aqueous pomegranate extract (AE-PG). Source: Gonzalez-Trujano et al. (2015); Cervantes-Anaya et al. (2022).

### 2.2. Animals and Housing

#### 2.2.1. Mice

Twenty-five adult BALB/c female mice (three months old) from the INPRFM vivarium were housed in groups of five at a room temperature of 22±2°C, under a 12/12-hour inverted light/dark cycle (light at 8 am), and a relative humidity of 50%, with food and water *ad libitum*. All experimental procedures were carried out in accordance with the guidelines established by the Institutional Ethics Committee for the Care and Use of Laboratory Animals CEI/CR/086/2015/2024 and the official Mexican Standard for the care and handling of animals (NOM-062-ZOO-1999). A technical assistant trained in animal welfare performed animal care and drug administration.

#### 2.2.2. Lambaris

Twenty-four adult lambaris (*Astyanax bimaculatus* Linnaeus 1758) were used in the present experiments. Animals were obtained from a colony resulting from cross-breeding between animals purchased from a local producer in Marabá/PA, Brazil. In the laboratory, the animals were acclimated for a minimum period of 15 days in 100-L water tanks. Tanks had constant mechanical aeration, biological filtration, artificial vegetation (De Sousa et al., 2024), and a 12-h light/12-h dark photoperiod. Feeding was provided twice daily with commercial feed (GUABI, Pirá line (28% crude protein, 5% fat, 300 mg/kg of vitamin C, and 87.5 IU/kg of vitamin E)) and captive-bred *Tenebrio molitor* larvae. Physicochemical parameters of the water were measured weekly in both tanks, averaging 0.8 ± 0.4 ppm ammonia, 6.5 ± 0.1 ppm dissolved oxygen, 1.1 ± 0.9 ppm nitrite, temperature of 26.7 ± 0.8 °C, pH 7 ± 0, conductivity (EC) 122 ± 8.5, and hardness of 60.0 ± 5.7 ppm. Brazilian standards of animal care for lambaris were followed (Conselho Nacional de Controle de Experimentação Animal - CONCEA, 2024; de Oliveira et al., 2023). The project was carried out with the approval of the Animal Use Ethics Committee of UNIFESSPA (Universidade Federal do Sul e do Sudeste do Pará) protocol number 23479.019576/2022-43.

#### 2.2.3. Zebrafish

Eighty-four adult zebrafish (*Danio rerio* Hamilton 1822) of the longfin phenotype were used in the present experiments. Animals were on average 4 months old when used in an experiment. Animals from both sexes were obtained from a colony resulting from cross-breeding between animals purchased in specialized stores (Power Fish Fisheries, Itaguaí/RJ, Brazil). Based on body morphology (general body shape and presence of genital papilla in females (Parichy et al., 2009)), a male:female ratio of approximately 40%:60% was estimated. All animals were kept in 50L tanks (density of 2 animals/L) and acclimatized in the laboratory for a period of 14 days before the onset of experiments. The tanks were kept constant at a temperature of 26 ± 2°C and pH 7.2, oxygenation, 14/10 light-dark cycle provided by fluorescent lamps, according to standards of care for zebrafish (Lawrence, 2007). Animals were fed twice a day with commercial flocked feed. Animal density adopted was in accordance with the protocol adopted in the Brazilian guideline for the care of zebrafish in the laboratory (Conselho Nacional de Controle de Experimentação Animal - CONCEA, 2024; de Oliveira et al., 2023). The project was carried out with the approval of the Animal Use Ethics Committee of UNIFESSPA (Universidade Federal do Sul e do Sudeste do Pará) protocol number 23479.019576/2022-43.

### 2.3. Stressors

#### 2.3.1. Acute Restraint Stress (Lambari and Zebrafish)

Zebrafish were assigned to two main groups: 24 that were maintained in control conditions, and 24 that were subjected to acute restraint stress (ARS). ARS protocols followed those described by Ghisleni et al. (2012), who showed that ARS for 90 min. produced both an increase in whole-body cortisol levels and a decrease in the expression of *crf* in the zebrafish brain. 30 min. before initiation of the protocol, Animals from the ARS group were submitted to acute restraint for 90 min. in plastic 1.5 mL microtubes with ∼5 mm openings in both extremities that allowed the circulation of water and oxygen. After lids were closed, the microtubes were inserted in a stand that was positioned in a 11 L tank filled with system water. Oxygenation was provided via an oxygen pump. Animals in the control group were placed in a tank similar to that of the stressed group, but without the restraint and rack, swimming freely in the aquarium for the same period of 90 min. The exposure was made under conditions of controlled temperature (23 °C ± 2), water pH (7.0 ± 1), lighting (549 ± 60 lux above the tank), and were made with the use of constant Gaussian white noise (62 dB). After stress, animals were evaluated in a behavioral battery with no interval between tests, composed of the novel tank test (NTT, 6 min) and the light/dark preference test (LDT, 15 min). The ARS protocol for lambaris was similar to that, with the exception that animals were restrained in 50 mL conical centrifuge tubes with ∼5 mm openings in both extremities, and animals the protocol was made in a 40 L tank; after stress, animals were evaluated in the NTT.

#### 2.3.2. Chronic Unpredictable Mild Stress (Mice)

Twenty-five female mice were assigned to two main groups: fourteen that were maintained under standard housing conditions in the institutional vivarium, and eleven that were subjected to unpredictable chronic mild stress (CUMS). For 21 days, CUMS mice were subjected to different stressors, while the control group (without CUMS) was manipulated daily in the vivarium. For the CUMS protocol, mice received two or three stressors per day, randomly applied to avoid adaptation (Vega-Rivera et al, 2014; Islas-Preciado et al., 2023). The stressor were: white noise, continuous light, dirty or wet cage, stroboscopic light, restricted movement, inclined cage, overcrowding (up to 10 animals per cage for 12 h) according to the protocol used by Vega-Rivera et al. 2014.

On day 14, both groups were subdivided into the following groups and treated for 7 days with a) the control group treated with saline solution (0.9%, i.p) and AE-PG (1.0 mg/kg, i.p). On day 22, 24 h after the last drug administration, all groups were evaluated in a behavioral battery with 15 min intervals between them, composed of open-field test (OFT, 5 min), elevated plus-maze (EPM, 5 min), splash test (ST, 5 min), and finally, forced swim test (FST, 6 min).

### 2.4. Behavioral assays

The behavioral assays used in the present work are the two most widely used tests in adult zebrafish, especially for anxiety-like behavior. While both tests use novelty as a stimulus to induce anxiety-like states (Kysil et al., 2017), the stimulus control differs: while the novel tank test (NTT) involves escape from the top, the light/dark test appears to involve an approach-avoidance conflict (Maximino et al., 2012). Thus, results from both tests are not always comparable. In all cases, animals were tested individually in each assay and were tested in a single assay.

#### 2.4.1. Fish

##### 2.4.1.1. Novel tank test

Both lambari and zebrafish were subjected to the novel tank test (NTT). While behavior in the NTT has been widely used as an index of anxiety-like behavior in zebrafish (Bencan et al., 2009; Egan et al., 2009; Levin et al., 2007), and described in *Astyanax mexicanus* (Padmanaban et al., 2025), it has not been used in *A. bimaculatus* yet. The novel tank test (NTT) assay was performed based on the protocol adapted from Cachat et al. (2010). The animals were transferred individually to a transparent tank measuring 20 cm x 25 cm x 15 cm (height x width x length) filled with 5 L of water. Upon transfer, the animals were allowed to explore the new aquarium freely for a period of 6 minutes, and this activity was recorded on an iPhone® (12 MP; 1080p resolution at 30 fps) positioned in front of the tank. Lighting and sound conditions were maintained throughout the experiment, with averages of 452 ± 40 lux and 66 ± 2 dB, respectively, under constant Gaussian white noise. Videos were analyzed using TheRealFishTracker software (v. 0.4.0; https://www.dgp.toronto.edu/~mccrae/projects/FishTracker/), focusing on behavior in the white compartment. The following endpoints were analyzed for all animals:

1. Geotaxis (s): Total time that the animal remained in the bottom third of the tank; higher geotaxis indicates an anxiogenic-like effect, while lower geotaxis indicates an anxiolytic-like effect (Bencan et al., 2009).
2. Erratic swimming (absolute turn angle): The absolute turn angle throughout the session; higher erratic swimming indicates an anxiogenic-like effect, while lower erratic swimming indicates an anxiolytic-like effect (Egan et al., 2009).
3. Average swimming speed (cm/s); changes in speed indicate hypo- or hyperlocomotion (Kalueff et al., 2013).

##### 2.4.1.2. Light/dark test (zebrafish)

In the light/dark preference test (LDT), a protocol adapted from Maximino et al. (2010) was adopted. Initially, zebrafish were transferred individually to the central compartment of a test tank measuring 15 cm x 10 cm x 45 cm (height x width x length). This aquarium had one white half and one black half, and the animals remained in the central compartment for 3 minutes to acclimatize. After this period, the partition separating the central compartment from the others was removed, allowing the animal to freely explore the other areas of the tank for 15 minutes. The animals’ activity during this interval was recorded on video using an iPhone® camera (12 MP; resolution of 1080p at 30 fps). Throughout the experiment, lighting conditions remained at an average of 472 ± 40 lux, and the average sound level was 67 ± 2 dB, with constant use of Gaussian white noise. Videos were analyzed using TheRealFishTracker software (v. 0.4.0; https://www.dgp.toronto.edu/~mccrae/projects/FishTracker/), focusing on behavior in the white compartment. The following endpoints were analyzed for each individual:

1. Scototaxis (s): Total time that the animal remained in the white compartment of the tank; higher scototaxis indicates an anxiogenic-like effect, while lower scototaxis indicates an anxiolytic-like effect (Maximino et al., 2010).
2. Erratic swimming (absolute turn angle): The absolute turn angle throughout the session; higher erratic swimming indicates an anxiogenic-like effect, while lower erratic swimming indicates an anxiolytic-like effect (Maximino et al., 2014).
3. Average swimming speed (cm/s); changes in speed indicate hypo- or hyperlocomotion (Kalueff et al., 2013).

#### 2.4.2. Mice

##### 2.4.2.1. Open field test

The animals were tested in an automated actimeter (Panlab, Harvard apparatus®, Mexico) in a 5 min session to discard the potential influence of treatments or stress conditions on general locomotor activity, as well as to understand the effects of the compounds in anxiety-like behavior. The apparatus consisted of a Plexiglas® cage (45 × 45 × 35 cm) with photo beams that detect in an automated manner: (a) time in the center of the apparatus; (b) total time in locomotion (horizontal activity); (c) frequency of stereotypical behavior; and (d) number of rearings (vertical exploration). The trajectories of animals were analyzed with an ActiTrack® system.

##### 2.4.2.2. Elevated plus-maze

To determine possible anxiolytic-like effects of treatments, mice were subjected to elevated plus-maze test (EPM) for mice. Briefly, EPM consisted of a two open arms (30 cm X 5 cm) perpendicular to two closed arms (30 cm X 5 cm X 16 cm) apparatus that was 40 cm above the floor. The apparatus contrasts two different zones: one was a potentially aversive zone (open arms), and the other was considered a safe zone (closed arms) (Walf & Frye, 2007). The animal was placed in the maze’s center, facing a closed arm. The session was video recorded (Canon PowerShot sx500 IS video-camera) for 5 min. The behavioral parameters registered were: (a) the cumulative time spent in the open arms, expressed as % of the time; (b) number of entries to open arms expressed as %; (c) the cumulative time in the closed arms, expressed as % of the time; (d) number of entries to open arms; and (e) the frequency of rearing behavior. Thus, more time in the open arms and more entries indicated less anxiety-like behavior; conversely, more time in the closed arms indicated a higher anxiety index (Walf & Frye, 2007). The total number of crosses was related to the general locomotor activity of the mice (Walf & Frye, 2007).

##### 2.4.2.3. Forced swim test

A modified forced swimming test (FST) was employed to assess the antidepressant-like effects of *P. granatum*. Mice were placed in a cylinder (30 cm height by 20 cm diameter) filled with water maintained at 21°C ± 1 for six minutes, and their behavior was recorded on video. An observer, blinded to treatment and group allocation, scored the recordings. During the final four minutes, the observer quantified the total time that the animal remains immobile and the total time spent in mobility behavior (swimming). Immobility was defined as minimal movement necessary to float or keep the head above water, while mobility was characterized by gentle movement around the cylinder (Flores-Gutiérrez et al 2018). A reduction in immobility behavior suggests of an antidepressant-like action (Porsolt et al., 1977).

##### 2.3.2.4. Splash test

Rodent grooming generally proceeds in a cephalocaudal direction (Kalueff et al., 2007). The behavioral sequence usually begins with licking of the paws, followed by washing the nose and face, the head, the body, the legs, and finally washing and licking the tail and genitals. Under stress conditions, the grooming pattern is altered and decreased, suggesting a deterioration of self-care, which is considered apathetic behavior, an index of depression (Kalueff et al., 2007). Antidepressant treatments increase grooming behavior.

After 30 min of acclimation, mice were subjected to the splash test in the two groups of mice. The test was conducted under dim light. An atomizer spray was used to splash a 10% sucrose solution twice on the dorsal coat of the mice, then quickly returned them to the home cage. After spraying the sucrose solution, the following variables were recorded: (a) latency to start grooming; (b) total grooming duration; (c) number of grooming bouts; (d) sequence of grooming patterns (pattern 1: elliptical stroke; pattern 2: unilateral stroke; pattern 3: bilateral stroke; pattern 4: body licking; Kalueff et al., 2007); (e) dominant grooming pattern (i.e., which grooming patterns was more frequent); (f) frequency of non-chain sequences (i.e., sequences in which the animal did not progress from pattern 1 to pattern 2, or pattern 2 to pattern 3, or pattern 3 to pattern 4, or pattern 4 to pattern 1); (g) frequency of chain sequences (i.e., sequences in which the animal did progress from pattern 1 to pattern 2, or pattern 2 to pattern 3, or pattern 3 to pattern 4, or pattern 4 to pattern 1).

### 2.5. Statistical Analysis

Two-way analyses of variance (ANOVAs) were applied to most behavioral data; when the assumptions of normality and/or homocedasticity were violated, Kruskal-Wallis two-way ANOVAs were used instead. For parametric ANOVAs, effect sizes are presented as partial eta-squared (η²p); for non-parametric ANOVAs, effect sizes are presented as epsilon-squared (ε²). Whenever p < 0.05, follow-up *post-hoc* tests were applied to the data, using Tukey’s HSD corrections for multiple comparisons. For data on dominant grooming patterns, multinomial logistic regressions were performed to test the effect of drug and stress on the dominant grooming pattern.

### 2.6. *In silico* Assays

#### 2.6.1. Molecule Selection, Target Prediction, and Network Pharmacology

Molecules from the phytochemical profile of the extract (Table 1) were analyzed in terms of drug-likeness in AdMetLab 3.0 (https://admetlab3.scbdd.com/). Molecules’ molecular weight (mw), predicted partition coefficient (logP), topological polar surface area (TPSA), quantitative estimate of drug-likeness (QED), plasma protein binding (PPB), and probability of crossing the blood-brain barrier (BBB) were calculated using the platform, and the Lipinski’s rule of five (which predicts good absorption of permeability), Pfizer’s rule (which predicts toxicity potential), and GSK rules (which predicts a favorable pharmacokinetic profile) were applied to these parameters to filter compounds with the highest likelihood of acting on the central nervous system. Table 2 presents these results; applying the three rules of thumb, the highest drug-likeness would be represented by ellagic acid (EA), with pelargonidin-3-O-glucoside (P3OG) in second place. These were retained for further analyses. While punicalagin was also found to produce antidepressant-like effects in ovariectomized rats (Cervantes-Anaya et al., 2022), its predicted poor absorption and low oral bioavailability and likely hydrolyzation into ellagic acid suggested that it is more likely a prodrug.

**Table 2.**
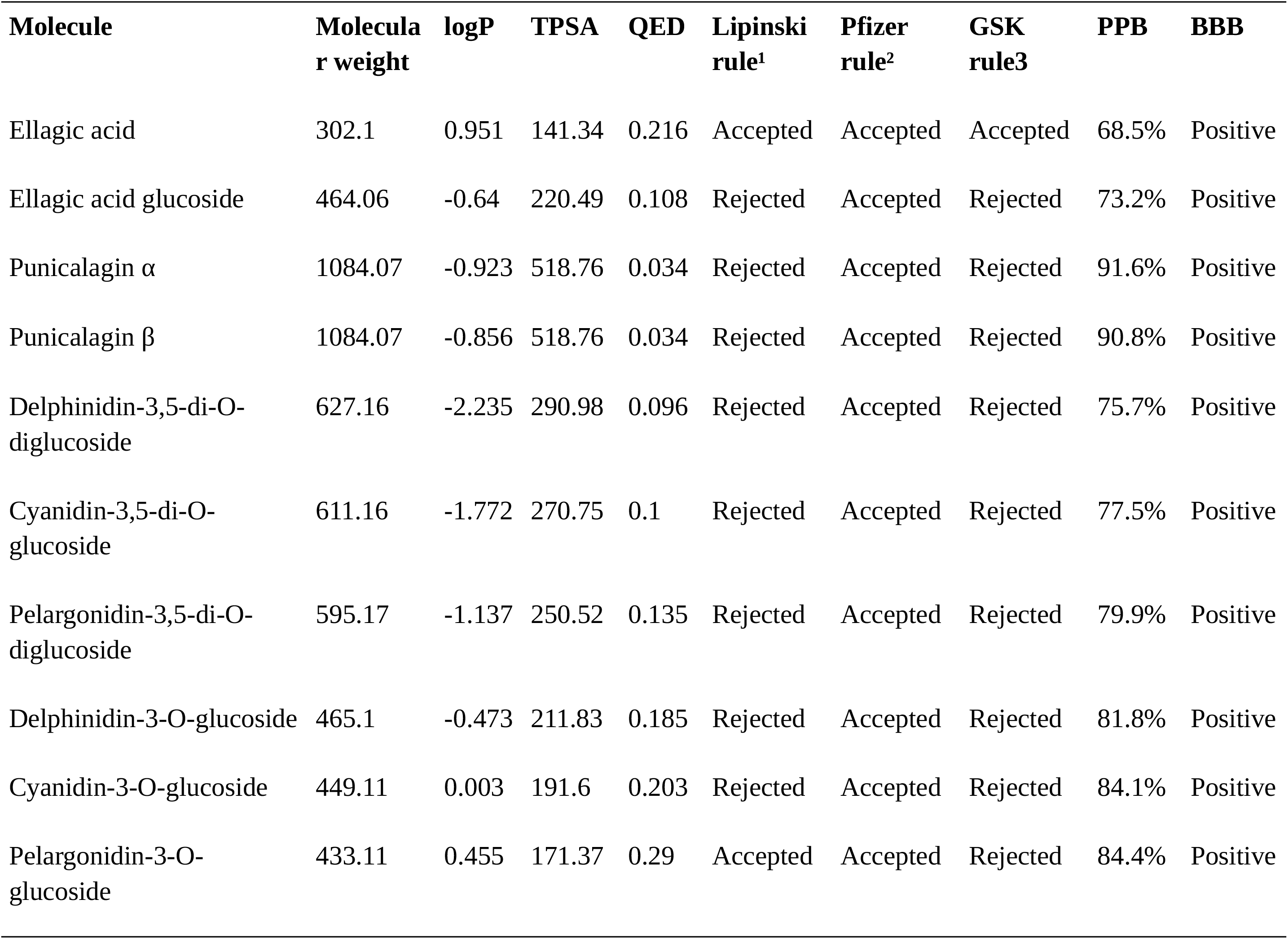
Predicted pharmacokinetic and toxicicity profile of the main polyphenolic compounds found in AE-PG.

| Molecule | Molecular weight | logP | TPSA | QED | Lipinski rule<sup>1</sup> | Pfizer rule<sup>2</sup> | GSK rule<sup>3</sup> | PPB | BBB |
| --- | --- | --- | --- | --- | --- | --- | --- | --- | --- |
| Ellagic acid | 302.1 | 0.951 | 141.34 | 0.216 | Accepted | Accepted | Accepted | 68.5% | Positive |
| Ellagic acid glucoside | 464.06 | -0.64 | 220.49 | 0.108 | Rejected | Accepted | Rejected | 73.2% | Positive |
| Punicalagin $\alpha$ | 1084.07 | -0.923 | 518.76 | 0.034 | Rejected | Accepted | Rejected | 91.6% | Positive |
| Punicalagin $\beta$ | 1084.07 | -0.856 | 518.76 | 0.034 | Rejected | Accepted | Rejected | 90.8% | Positive |
| Delphinidin-3,5-di-O-diglucoside | 627.16 | -2.235 | 290.98 | 0.096 | Rejected | Accepted | Rejected | 75.7% | Positive |
| Cyanidin-3,5-di-O-glucoside | 611.16 | -1.772 | 270.75 | 0.1 | Rejected | Accepted | Rejected | 77.5% | Positive |
| Pelargonidin-3,5-di-O-diglucoside | 595.17 | -1.137 | 250.52 | 0.135 | Rejected | Accepted | Rejected | 79.9% | Positive |
| Delphinidin-3-O-glucoside | 465.1 | -0.473 | 211.83 | 0.185 | Rejected | Accepted | Rejected | 81.8% | Positive |
| Cyanidin-3-O-glucoside | 449.11 | 0.003 | 191.6 | 0.203 | Rejected | Accepted | Rejected | 84.1% | Positive |
| Pelargonidin-3-O-glucoside | 433.11 | 0.455 | 171.37 | 0.29 | Accepted | Accepted | Rejected | 84.4% | Positive |

Probable targets of the selected compounds (ellagic acid and pelargonidin-3-O-glucoside) were searched for in the Traditional Chinese Medicine Database (https://tcmsp-e.com/tcmsp.php) and Comparative Toxicogenomics Database (https://ctdbase.org/search/); their SMILES notations were also input on SWISS TargetPrediction (https://www.swisstargetprediction.ch/). A total of 186 targets were found for ellagic acid, and 36 for pelargonidin-3-*O*-glucoside (P3OG). Targets involved in anxiety were retrieved from GeneCards (https://www.genecards.org/search/results?q=anxiety); a total of 2540 targets were found as anxiety-related proteins. To construct the drug-target network, we first used jVenn (Bardou et al., 2014) (https://www.bioinformatics.com.cn/static/others/jvenn/example.html) to identify the intersection of predicted compound targets and proteins involved in anxiety. This produced a total of 11 proteins (Figure 1). For these intersection targets, the STRING database (v. 12.0; https://stringdb.org) was used to construct a protein-protein interaction (PPI) network of common targets. The network was configured to filter no less than 20 interactors in the second shell, with the first shell consisting of only the queried proteins; to enter the network, a given interaction needed to have a minimum interaction score of 0.9, representing high confidence. The tsv files were downloaded and imported into Cytoscape (v. 3.10.4)(Shannon et al., 2003) for visualization and further analysis. The network topology was analyzed using degree centrality parameters. To further understand the role of the targets, we identified pathways by enrichment analysis using the Reactome pathway database (https://reactome.org/PathwayBrowser/) inside Cytoscape.

**Figure 1.**
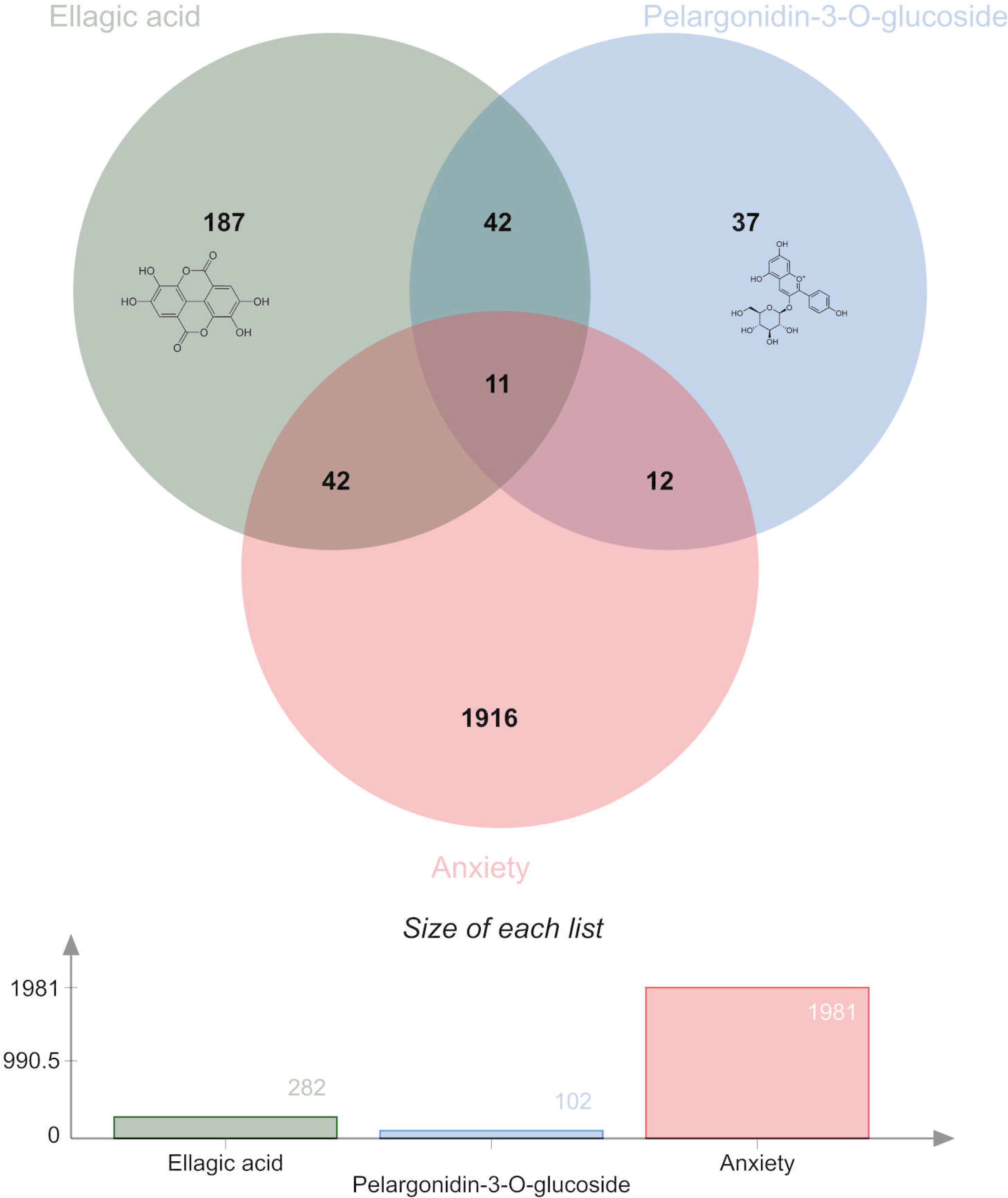
Venn diagram of proteins involved in anxiety disorders and targets shared by ellagic acid and pelargonidin-3-O-glucoside.

#### 2.6.2. Predicted Expression Analysis

##### Data set

Regional expression data for 39 genes across 12 limbic-associated brain regions (long insular gyri [LIG], short insular gyri [SIG], anterior cingulate gyrus [CgGf], basolateral amygdala [BLA], central amygdala [CeA], septal nuclei [SptN], habenula [Hb], paraventricular thalamus [Pa], periventricular hypothalamic nucleus [PVH], central gray of midbrain [CGMB], midbrain raphe [MBRa], and locus coeruleus [LC]) were obtained from the Allen Brain Atlas (Allen Institute for Brain Science, 2025) and supplied as relative expression levels (z-scores). Because the raw values span two orders of magnitude, the matrix was normalized prior to clustering, as implemented in Cluster 3.0 (de Hoon et al., 2004). The normalized matrix is symmetric around zero and compressed to a range of approximately ±0.5.

##### Hierarchical clustering

Genes and arrays were clustered independently in Cluster 3.0 using Pearson correlation as the similarity measure and centroid linkage. The resulting trees were visualized in TreeView (Saldanha, 2004). Gene clusters were defined by cutting the gene dendrogram at *k* = 2 and *k* = 4, and regional clusters by cutting the array dendrogram at *k* = 3.

##### Principal component analysis (PCA)

PCA was performed in Cluster 3.0 (de Hoon et al., 2004) on the same normalized matrix, with genes and arrays analyzed in a shared subspace, yielding paired eigenvector scores and region and gene coordinates outputs for the first 11 components. Eigenvalues were extracted and expressed as percentages of the summed eigenvalues. The two PCA runs (array-PCA and gene-PCA) were cross-checked for numerical consistency; eigenvalues agree to within ∼1%, confirming a valid dual decomposition.

#### 2.6.3 Molecular Docking

Since SWISS TargetPrediction predicted that both EA and P3OG would interact with ERα/ERS1 and ERβ/ERS2, and since the effects of AE-PG were shown to be mediated by the ERβ receptor in ovariectomized rats (Valdés-Sustaita et al., 2021), we performed molecular interaction simulations between these two compounds and the binding domain of ERβ in SwissDock (https://www.swissdock.ch/). The structure coordinates of EA (PubChem CID: 5487461) and P3OG (PubChem CID: 443648) were obtained from the PubChem database (https://pubchem.ncbi.nlm.nih.gov/) in SMILES format. The initial coordinates of the target human estrogen receptor β ligand-binding domain (PDB: 1QKM) were obtained from the RCSB PDB database (https://www.rcsb.org/). SwissDock (Bugnon et al., 2024) was used to prepare the ligand and the target for docking. The target was optimized in SwissDock using AutoDock Vina (Eberhardt et al., 2021) to remove all water molecules and the genistein ligand in the original structure and adding Kollman charges and nonpolar hydrogens. AutoDock Vina was used for the preparation of the grid box on SwissDock. A search space was prepared using a 13 × 13 × 13 Å (X = 23.0, Y = 8.0, Z = 114.0), coinciding with the region in which genistein was shown to interact with the protein (Pike et al., 1999). Molecular docking as performed using the SwissDock webserver, using AutoDock Vina to calculate an affinity constant (Bugnon et al., 2024; Eberhardt et al., 2021).

## 3. Results

### 3.1. Behavioral Assays

#### 3.1.1. Lambari NTT

In *Astyanax bimaculatus (*Figure 2A), a large main effect of stress was observed for geotaxis (F_[1, 19]_ = 8.301, p = 0.01, η²p = 0.304), but the medium-sized effect of treatment was not statistically significant (F_[1, 19]_ = 1.769, p = 0.199, η²p = 0.116). A significant and large interaction effect was found for this main endpoint (F_[1, 19]_ = 4.722, p = 0.043, η²p = 0.238; Figure 2B). Post-hoc tests found that stress increased time on bottom in untreated animals (S^-^ + 0 mg/kg vs. ARS + 0 mg/kg, *d* = -2.196, p = 0.009), an effect that was partially blocked by AE-PG (ARS + 0 mg/kg vs. ARS + 1.0 mg/kg, *d* = 1.681, p = 0.05).

**Figure 2.**
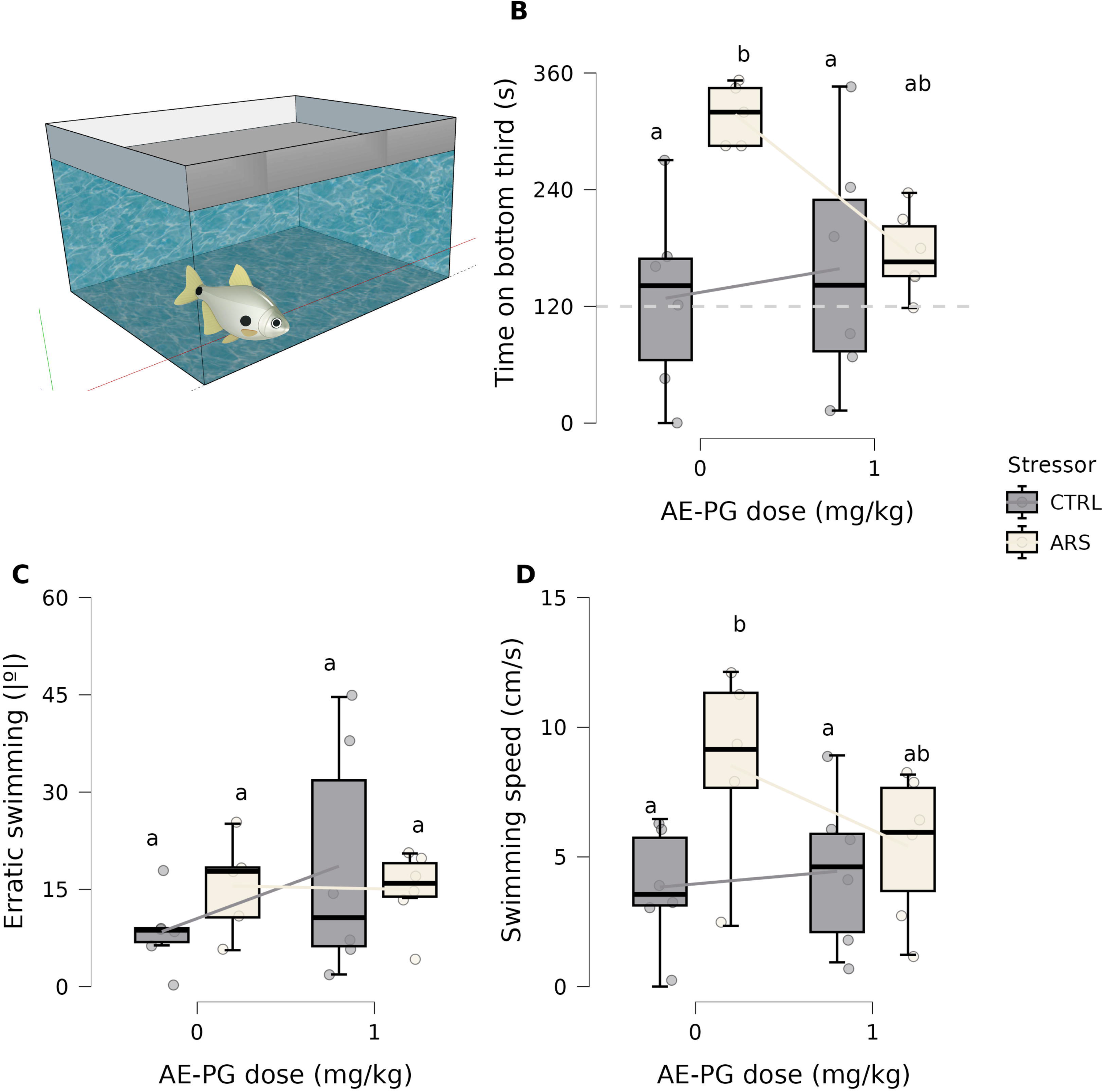
Behavioral effects of AE-PG (1 mg/kg) on lambari (*A. bimaculatus*) previously exposed to acute restraint stress (ARS) and tested in the novel tank test. (A) Schematics of the test tank. (B) Geotaxis; the dotted line represents time spent at the bottom of the tank at the chance level. (C) Erratic swimming. (D) Swimming speed. Dots represent data from individual animals. Letters above each boxplot represent statistically significant differences at the p < 0.05 level; different letters represent significant differences, while groups with equal letters do not differ at the p < 0.05 level. Boxplots represent median ± interquartile range; the lines between boxes connect group averages.

No main effects were found for erratic swimming (Figure 2C; stress: F_[1, 19]_ = 0.0.146, p = 0.707, η²p = 0.008; treatment: F_[1, 19]_ = 1.125, p = 0.302, η²p = 0.056); an interaction effect was also absent (F_[1, 19]_ = 1.385, p = 0.254, η²p = 0.067). Finally, a medium main effect of stress was found for swimming speed (Figure 2D; F_[1, 19]_ = 5.016, p = 0.037, η²p = 0.208), but a main effect of treatment was absent (F_[1, 19]_ = 0.9871, p = 0.333, η²p = 0.049), and the medium interaction effect was also not statistically significant (F_[1, 19]_ = 2.171, p = 0.157, η²p = 0.103). Post-hoc tests found that ARS increased swimming speed in lambari (Figure 2D; *d* = 0.934, p = 0.037).

#### 3.1.2. Zebrafish NTT

In the novel tank test in zebrafish (Figure 3A), a large main effect of stress was found for geotaxis (F_[1, 44]_ = 12.15, p = 0.001, η²p = 0.216), and a medium main effect of treatment was observed (F _[1, 44]_ = 4.998, p = 0.03, η²p = 0.102); a large interaction effect was also found (F_[1, 44]_ = 15.51, p < 0.001, η²p = 0.261). Post-hoc tests showed that ARS significantly increased geotaxis (Figure 3B, d = -2.143, p < 0.001), an effect that AE-PG prevented (ARS + 0 mg/kg vs. ARS + 1.0 mg/kg, *d* = 1.782, p < 0.001). Similar effects were observed on erratic swimming (stress: F_[1, 43]_ = 15.0, p < 0.001, η²p = 0.259; treatment: F_[1, 43]_ = 30.08, p < 0.001, η²p = 0.412; interaction: F_[1, 43]_ = 10.68, p = 0.002, η²p = 0.199). Post-hoc tests found that ARS significantly increased erratic swimming (Figure 3C, *d* = -2.085, p < 0.001), and AE-PG prevented this effect (ARS + 0 mg/kg vs. ARS + 1.0 mg/kg, *d* = 2.555, p < 0.001). Finally, no effects were observed on swimming speed (Figure 3; stress: F_[1, 44]_ = 0.04, p = 0.842, η²p = 9.1 x 10^-4^; dose: F_[1, 44]_ = 1.049, p = 0.311, η²p = 0.023; interaction: F_[1, 44]_ = 0.223, p = 0.639, η²p = 0.005).

**Figure 3.**
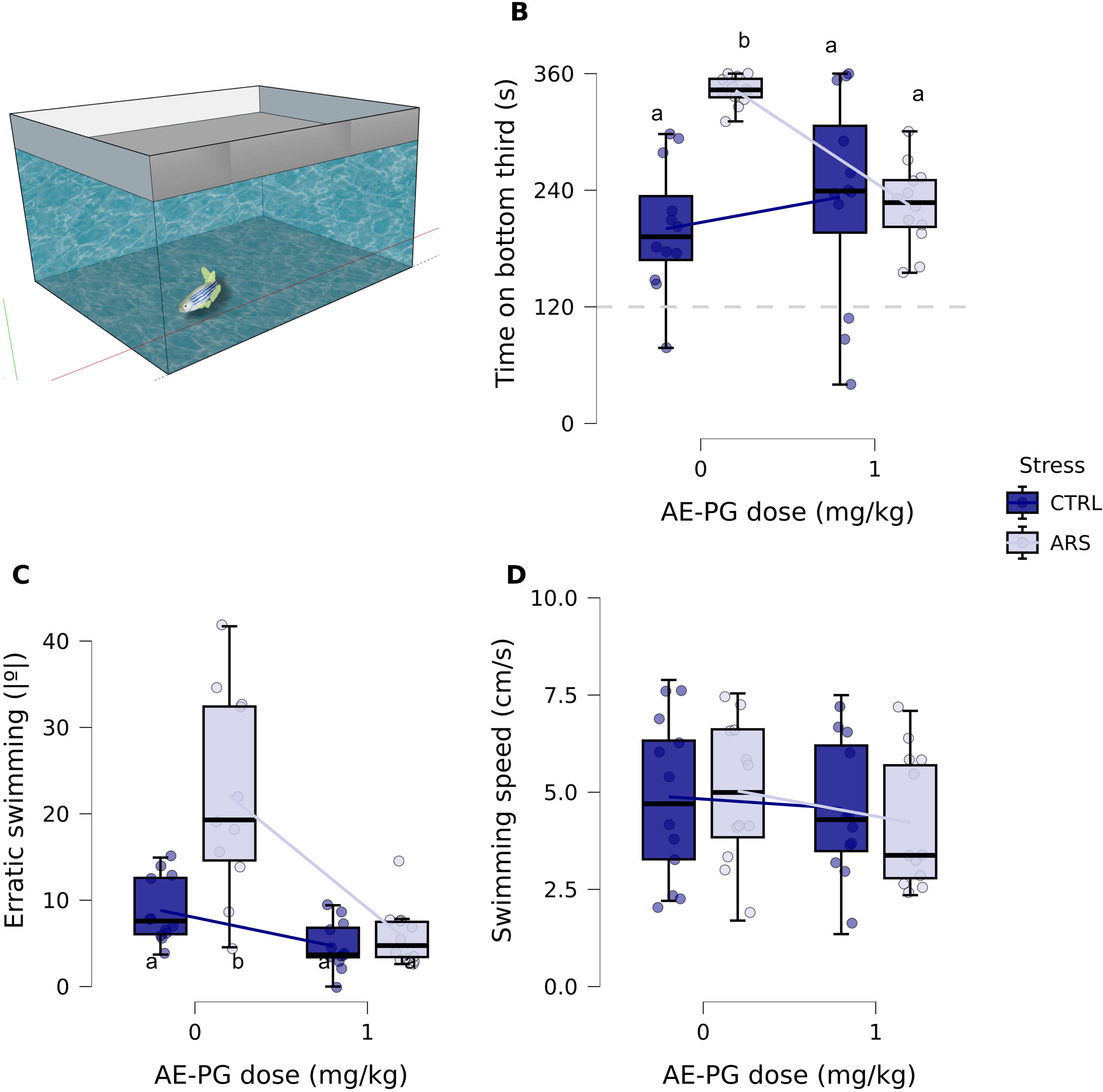
Behavioral effects of AE-PG (1 mg/kg) on zebrafish (*D. rerio*) previously exposed to acute restraint stress (ARS) and tested in the novel tank test. (A) Schematics of the test tank. (B) Geotaxis; the dotted line represents time spent at the bottom of the tank at the chance level. (C) Erratic swimming. (D) Swimming speed. Dots represent data from individual animals. Letters above each boxplot represent statistically significant differences at the p < 0.05 level; different letters represent significant differences, while groups with equal letters do not differ at the p < 0.05 level. Boxplots represent median ± interquartile range; the lines between boxes connect group averages.

#### 3.1.3. Zebrafish LDT

In the light dark test in zebrafish (Figure 4A), no main effect of stress was found for the main endpoint, time on white (F_[1, 43]_ = 0.411, p = 0.525, η²p = 0.009), but a large main effect of treatment was found (F_[1, 43]_ = 30.93, p < 0.001, η²p = 0.418); a large interaction effect was also found for this variable (F_[1, 43]_ = 13.23, p < 0.001, η²p = 0.235). Post-hoc tests found that ARS increased scototaxis (i.e., decreased time on white; *d* = 1.249, p = 0.023), an effect that was blocked by AE-PG (ARS + 0 mg/kg vs. ARS + 1.0 mg/kg: *d* = -2.685, p < 0.001)(Figure 4B). No main effect of stress was found for risk assessment (F_[1, 43]_ = 1.824, p = 0.184, η²p = 0.041), but a large main effect of treatment was found for this endpoint (F_[1, 43]_ = 16.71, p < 0.001, η²p = 0.279). A medium-sized interaction effect was found for risk assessment (F_[1, 43]_ = 6.936, p = 0.012, η²p = 0.139). Post-hoc tests suggested that ARS increased erratic swimming (*d* = -1.163, p = 0.038), an effect prevented by AE-PG (ARS + 0 mg/kg vs. ARS + 1.0 mg/kg: *d* = 1.962, p < 0.001)(Figure 4C). No effects were observed for thigmotaxis (stress: F_[1, 43]_ = 0.018, p = 0.892, η²p = 4.305 x 10^-4^; dose: F_[1, 43]_ = 0.024, p = 0.876, η²p = 5.683 x 10^-4^; interaction: F_[1, 43]_ = 3.785, p = 0.058, η²p = 0.081)(Figure 4D). A medium main effect of stress was found for erratic swimming (F _[1, 43]_ = 5.879, p = 0.02, η²p = 0.12), but no significant effect of treatment was found (F_[1, 43]_ = 2.735, p = 0.105, η²p = 0.059); interactions were also absent (F_[1, 43]_ = 1.336, p = 0.254, η²p = 0.03); ARS increased erratic swimming regardless of treatment (*d* = -0.708, p = 0.02)(Figure 4E). No effects were observed for freezing (stress: F_[1, 43]_ = 2.62, p = 0.113, η²p = 0.057; dose: F_[1, 43]_ = 0.537, p = 0.468, η²p = 0.012; interaction: F_[1, 43]_ = 0.158, p = 0.693, η²p = 0.004)(Figure 4F). A large main effect of stress was found for entries on white (F_[1, 43]_ = 43.2, p < 0.001, η²p = 0.501), but treatment (F_[1, 43]_ = 1.676, p = 0.202, η²p = 0.037) and interaction (F_1[, 43]_ = 1.19, p = 0.281, η²p = 0.027) effects were not statistically significant (Figure 4G); ARS decreased entries on white regardless of AE-PG treatment (*d* =1.919, p < 0.001). Finally, no effects were observed on swimming speed (Figure 4H; stress: F_[1, 43]_ = 0.121, p = 0.73, η²p = 0.003; dose: F_[1, 43]_ = 0.034, p = 0.855, η²p = 7.8 x 10^-4^; interaction: F_[1, 43]_ = 1.021, p = 0.224, η²p = 0.034).

**Figure 4.**
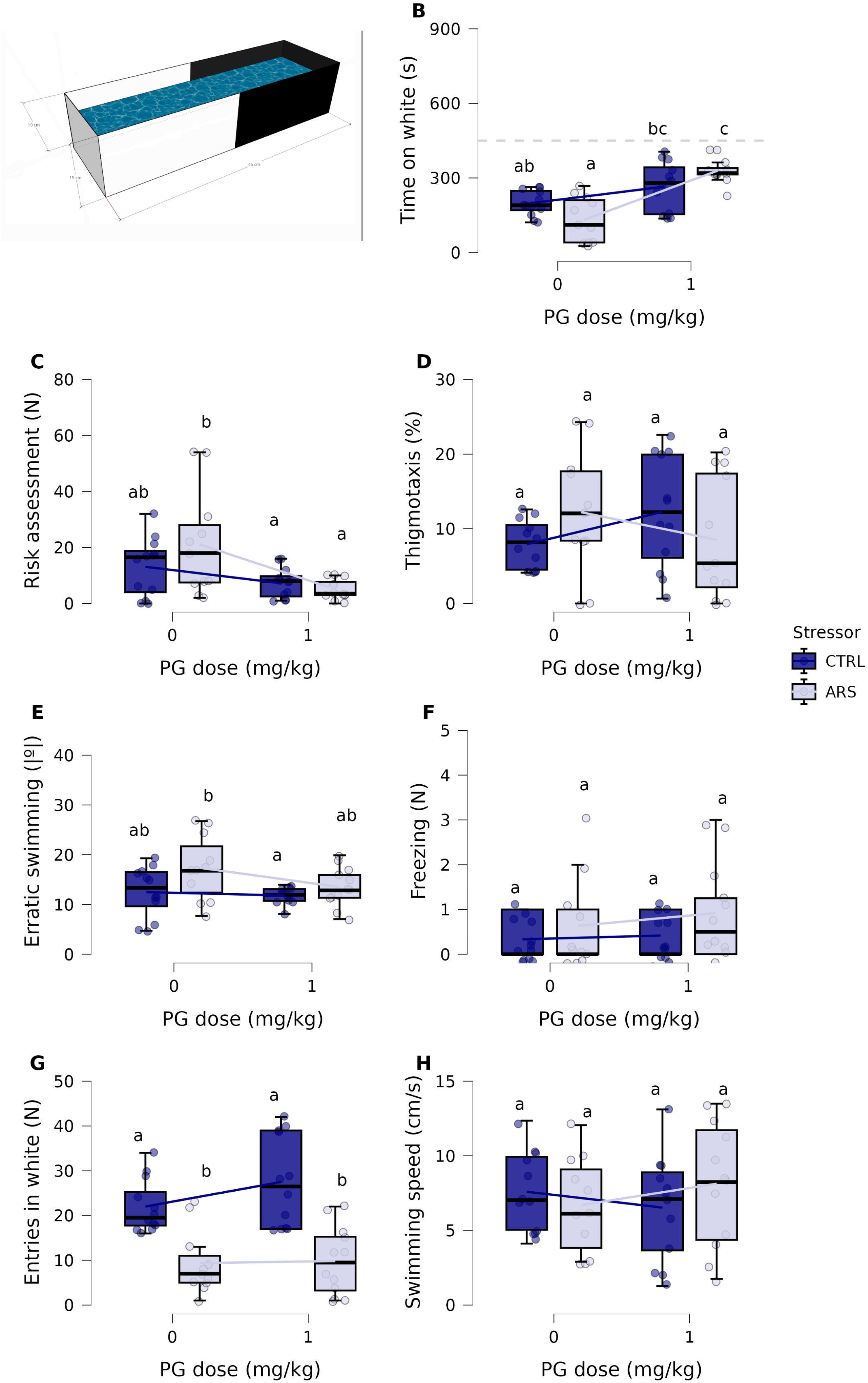
Behavioral effects of AE-PG (1 mg/kg) on zebrafish (*D. rerio*) previously exposed to acute restraint stress (ARS) and tested in the light/dark test. (A) Schematics of the test tank. (B) Scototaxis; the dotted line represents time spent at the white side of the tank at the chance level. (C) Risk assessment. (D) Thigmotaxis. (E) Erratic swimming. (F) Freezing. (G) Entries in the white compartment. (H) Swimming speed. Dots represent data from individual animals. Letters above each boxplot represent statistically significant differences at the p < 0.05 level; different letters represent significant differences, while groups with equal letters do not differ at the p < 0.05 level. Boxplots represent median ± interquartile range; the lines between boxes connect group averages.

#### 3.1.4. Mouse EPM

In the mouse elevated plus-maze (Figure 5A), no significant main effect of stress was found for percentage of time in the open arms (F_[1, 21]_ = 2.301, p = 0.144, η²p = 0.099), but a large main effect of treatment was found (F_[1, 21]_ = 5.09, p = 0.035, η²p = 0.195); a significant interaction effect was also found (F_[1, 21]_ = 6.204, p = 0.021, η²p = 0.228)(Figure 5B). *Post-hoc* tests found that CUMS decreased percentage of time in the open arms (CTRL + 0 mg/kg vs. CUMS + 0 mg/kg: t_[df = 21]_ = 2.764, p = 0.05, *d* = 1.619), and AE-PG reversed this effect (CUMS + 0 mg/kg vs. CUMS + 1 mg/kg: t_[df = 21]_ = t_[df = 21]_ = -3.166, p = 0.022, *d* = -1.917); AE-PG itself had no effect on non-stressed animals (CTRL + 0 mg/kg vs. CTRL + 1 mg/kg: t_[df = 21]_ = 0.1773, p = 0.998, *d* = 0.095). No significant main effect of stress was found for entries in the open arms (H_[df = 1]_ = 0.814, p = 0.367, ε² = 0.034), but a significant and large main effect of treatment was found (H_[df = 1]_ = 7.234, p = 0.007, ε² = 0.301)(Figure 5C). *Post-hoc* tests found that treatment with 1 mg/kg AE-PG showed more entries in the open arms irrespective of stress (z = -2.69, p = 0.007). No significant main effect of stress (F_[1. 21]_ = 3.116, p = 0.092, η²p = 0.123) nor treatment (F_[1, 21]_ = 4.078, p = 0.056, η²p = 0.163) were found for percentage of time in the closed arms; an interaction effect was also absent (F_[1, 21]_ = 1.502, p = 0.234, η²p = 0.067)(Figure 5D). No significant main effects of stress (H_[df = 1]_ = 0.092, p = 0.762) or treatment (H_[df = 1]_ = 2.348, p= 0.125) were found for total entries (Figure 5E). Finally, no significant main effects of stress (F_[1, 21]_ = 0.61, p = 0.444, η²p = 0.031) or treamtent (F_[1, 21]_ = 0.447, p = 0.511, η²p = 0.021) were found for rearings (5F), but a large interaction effect was found (F_[1, 21]_ = 6.01, p = 0.023, η²p = 0.222); however, after *post-hoc* tests no differences were found between groups.

**Figure 5.**
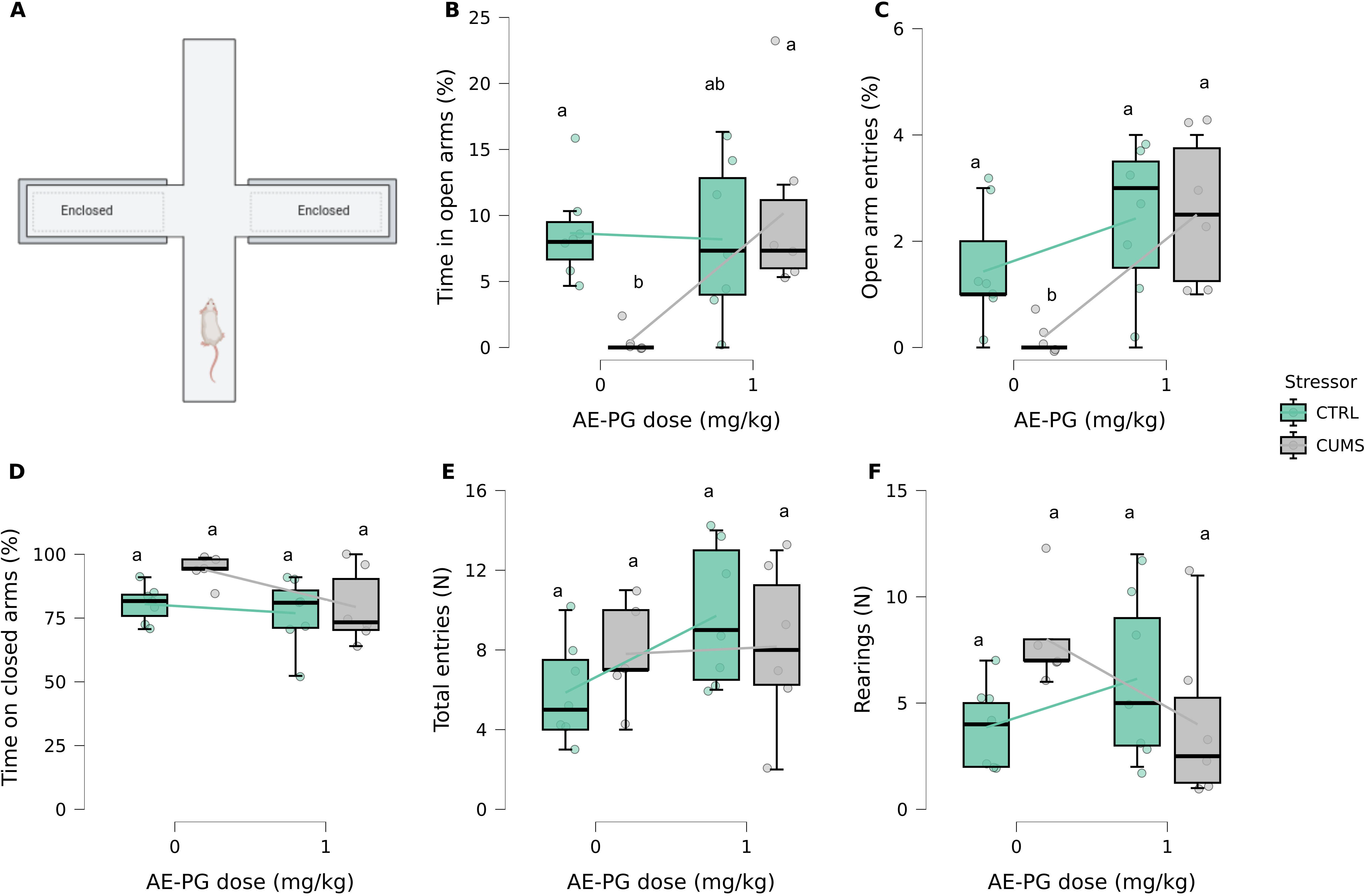
Behavioral effects of AE-PG (1 mg/kg) on mice (*M. musculus*) previously exposed to chronic unpredictable mild stress (CUMS) and tested in the elevated plus-maze. (A) Schematics of the test apparatus. (B) Percentage of time spent in the open arms. (C) Percentage of number of entries in the open arms. (D) Percentage of time in the closed arms. (E) Total entries in any arm. (F) Frequency of rearings. Dots represent data from individual animals. Letters above each boxplot represent statistically significant differences at the p < 0.05 level; different letters represent significant differences, while groups with equal letters do not differ at the p < 0.05 level. Boxplots represent median ± interquartile range; the lines between boxes connect group averages.

#### 3.1.5. Mouse open-field test

In the open field test (Figure 6A), a significant main effect of stress was found for center time (F_[1, 21]_ = 4.863, p = 0.039, η²p = 0.188), but no main effects of treatment were found (F_[1, 21]_ = 0.3926, p = 0.538, η²p = 0.018); interaction effects were also not found (F_[1, 21]_ = 0.642, p = 0.432, η²p = 0.029). *Post-hoc* tests found that CUMS decreased center time in both stressed and unstressed animals (t_[df = 21]_ = 2.205, p = 0.039, *d* = 0.891)(Figure 6B). A significant main effect of stress was found for total time in locomotion (F_[1, 21]_ = 5.649, p = 0.027, η²p = 0.212), but no effects of treatment (F_[1, 21]_ = 4.25 x 10^-4^, p = 0.984, η²p = 2.024 x 10^-5^) or interaction (F_[1, 21]_ = 0.452, p = 0.509, η²p = 0.021) were found for this variable; *post-hoc* tests revealed that CUMS decreased time in locomotion in both unstressed and stressed animals (t_[df = 21]_ = 2.377, p = 0.027, *d* = 0.9599) (Figure 6C). No main effects of stress or treatment were found for frequency of stereotypy (stress: H_[df = 1]_ = 1.589, p = 0.683, ε² = 0.066; dose: H_[df = 1]_ = 0.167, p 0.683, ε² = 0.007)(Figure 6D). Finally, a significant main effect of stress (H_[df = 1]_ = 10.45, p 0.001, ε² = 0.435), but not treatment (H_[df = 1]_ = 0.96, p = 0.327, ε² = 0.04), was found for frequency of rearing; *post-hoc* tests revealed that CUMS decreased rearing irrespective of AE-PG treatment (z = 3.232, p = 0.001)(Figure 6E).

**Figure 6.**
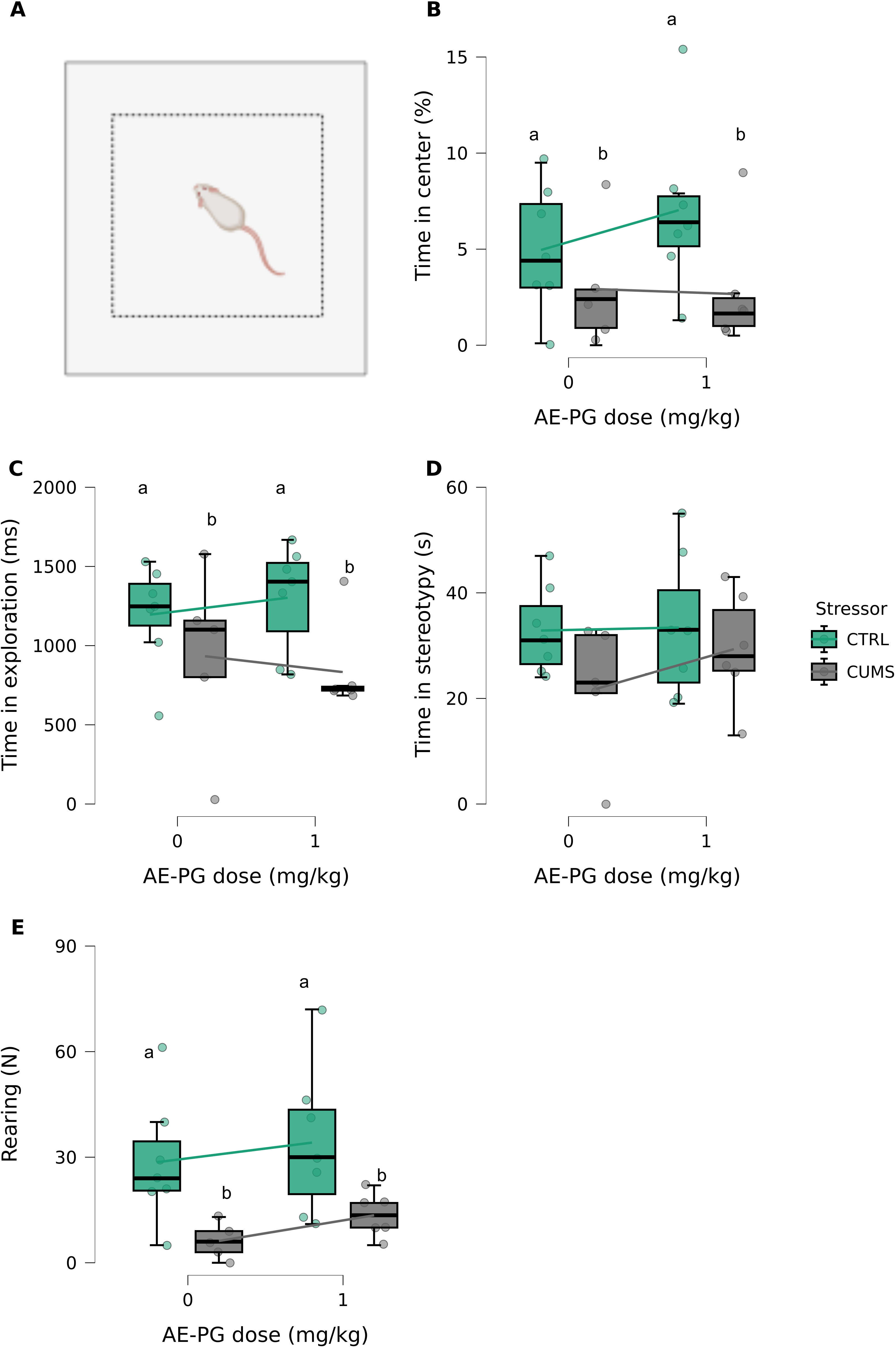
Behavioral effects of AE-PG (1 mg/kg) on mice (*Balb-c*) previously exposed to chronic unpredictable mild stress (CUMS) and tested in the open field-test (OFT). (A) Schematics of the test apparatus. (B) Percentage of time spent in the center of the apparatus. (C) Total time exploring the apparatus. (D) Total time displaying stereotypical behavior. (E) Frequency of rearings. Dots represent data from individual animals. Letters above each boxplot represent statistically significant differences at the p < 0.05 level; different letters represent significant differences, while groups with equal letters do not differ at the p < 0.05 level. Boxplots represent median ± interquartile range; the lines between boxes connect group averages.

#### 3.1.6. Mice splash test

In the mice splash test, no significant main effects of stress (F_[1, 21]_ = 0.278, p = 0.603, η²p = 0.013) or treatment (F_[1, 21]_ = 1.975, p = 0.175, η²p = 0.086) were found for grooming latency; no interaction effect was also found (F_[1, 21]_ = 0.408, p = 0.53, η²p = 0.019)(Figure 7A). A significant and large main effect of stress was found for total grooming duration (F_[1, 21]_ = 7.224, p = 0.014, η²p = 0.256), but no significant effect of treatment was found (F_[1, 21]_ = 1.48 x 10^-4^, p = 0.99, η²p = 70.2 x 10^-6^); interaction effects were also absent (F_[1, 21]_ = 2.171, p = 0.155, η²p = 0.094). *Post-hoc* tests found that CUMS decreased grooming duration (CTRL + 0 mg/kg vs. CUMS + 0 mg/kg: t_[df = 21]_ = 2.87, p = 0.042, *d* = 1.68), but AE-PG did not rescue this effect (CUMS + 0 mg/kg vs. CUMS + 1 mg/kg: t_[df = 21]_ = -0.991, p = 0.756, *d* = -0.6), nor did it have an effect by itself (CTRL + 0 mg/kg vs. CTRL + 1 mg/kg: t_[df = 21]_ = 1.104, p = 0.691, *d* = 0.59)(Figure 7B). No main effect of stress was found for total grooming bouts (H_[df = 1]_ = 0.597, p = 0.44, ε² = 0.025), but a significant effect of treatment was found (H_[df = 1]_ = 9.607, p = 0.002, ε² = 0.4); *post-hoc* tests revealed that treatment with AE-PG decreased the frequency of grooming bouts irrespective of stress exposure (z = 3.1, p = 0.002)(Figure 7C). Qualitative analysis of the syntactic pattern of grooming in the splash test (Figure 7D) suggests that CUMS disrupts the normal sequence (cephalocaudal progression), with more transitions between unilateral stroke and body licking, fewer transitions between unilateral and bilateral stroke, and no occurrence of elliptical strokes; however, AE-PG also seemed to disrupt the grooming sequence, with no elliptical strokes and no sequence of bilateral strokes in non-stressed animals treated with the extract, and no body licking in stressed animals treated with the extract. A multinomial logistic regression was performed to test whether the effect of treatment on the distribution of dominant grooming patterns (1–4) was moderated by stress exposure; the overall interaction between treatment and stress was not statistically significant (χ²_[df - 63]_) = 13.84, p = 0.128)(Figure 7E). However, the small sample size and sparse distribution across the 4 grooming categories limit the statistical power of this analysis, and these null findings should be interpreted cautiously. No main effect of stress (H_[df = 1]_ = 0.028, p = 0;868, ε² = 0.001) was found for the frequency of non-chain sequences, but a main effect of treatment was found (H_[df = 1]_ = 11.67, p < 0.001); *post-hoc* tests revealed that treatment with AE-PG reduced the frequency of non-chain sequences (z = 3.416, p < 0.001)(Figure 7F). Finally, a main effect of stress (H_[df = 1]_ = 7.735, p = 0.005, ε² = 0.322) was found for the frequency of chain sequences, but a main effect of treatment was absent (H_[df = 1]_ = 0.024, p = 0.877, ε² = 0.001); *post-hoc* tests found that CUMS decreased the frequency of chain sequences (z = 2.781, p = 0.005)(Figure 7G).

**Figure 7.**
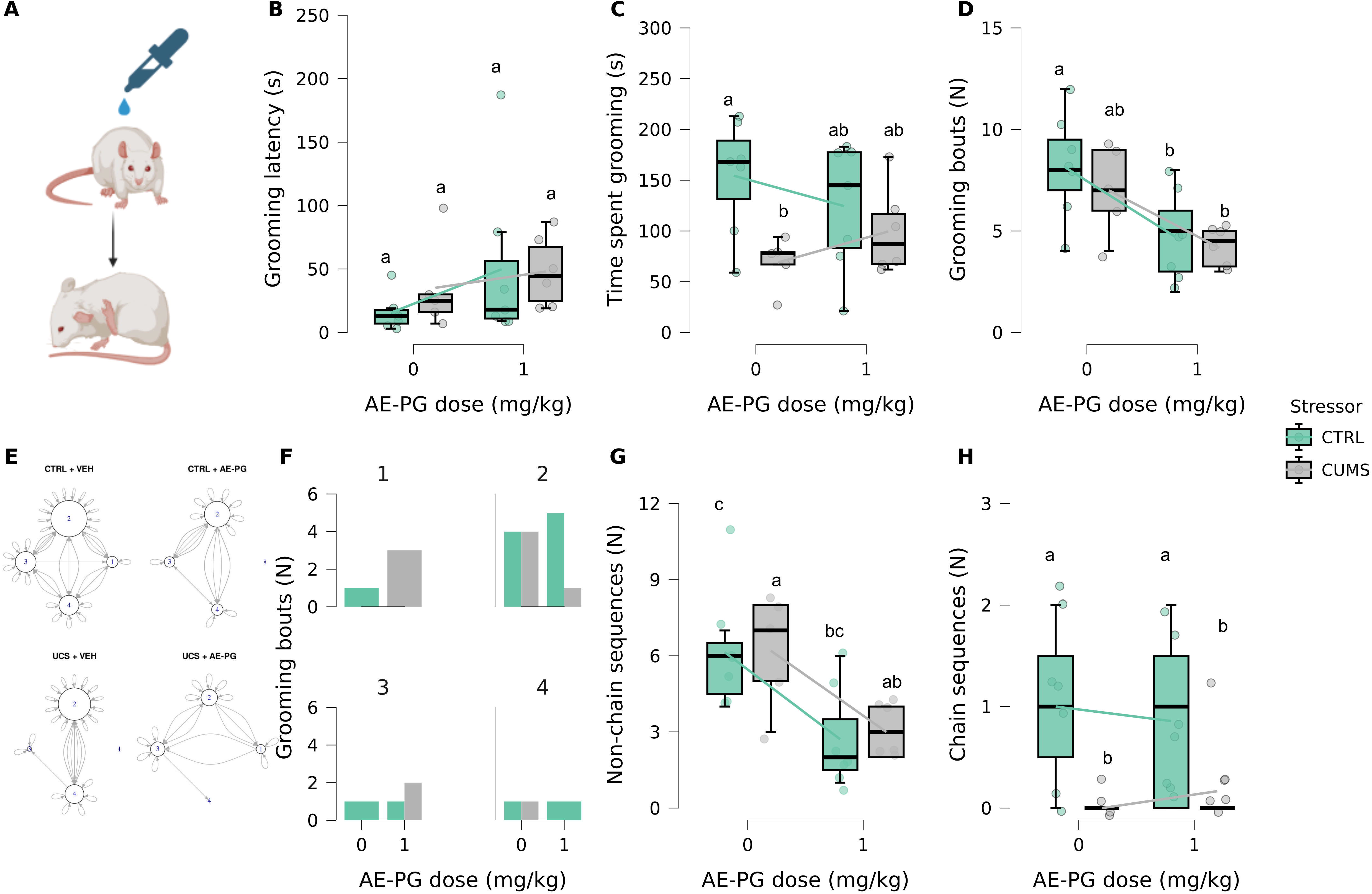
Behavioral effects of AE-PG (1 mg/kg) on mice (Balb-c) previously exposed to unpredictable chronic stress (CUMS) and tested in the splash. (A) Schematics of the test. (B) Latency to start grooming. (C) Total time spent grooming. (D) Number of grooming bouts. (E) Behavioral transitions between grooming patterns. (F) Number of grooming bouts per chain element. (G) Frequency of non-chain sequences. (H) Frequency of chain sequences. Dots represent data from individual animals. Letters above each boxplot represent statistically significant differences at the p < 0.05 level; different letters represent significant differences, while groups with equal letters do not differ at the p < 0.05 level. Boxplots represent median ± interquartile range; the lines between boxes connect group averages.

#### 3.1.7. Mice FST

In the forced swim test, no significant main effect of stress was found for total immobility time (F_[1, 21]_ = 1.393, p = 0.215, η²p = 0.062), but a significant main effect of treatment was found (F_[1, 21]_ = 47.76, p < 0.001, η²p = 0.695); interaction effects were not found (F_[1, 21]_ = 2.425, p = 0.134,η²p = 0.103). *Post-hoc* tests found that AE-PG decreased immobility time in both stressed and unstressed animals (t_[df = 21]_ = 6.911, p < 0.001, *d* = 2.791)(Figure 8A). No main effect of stress was found for total time in activity (F_[1, 21]_ = 1.412, p = 0.248, η²p = 0.063), but a significant main effect of treatment was found (F_[1, 21]_ = 47.65, p < 0.001, η²p = 0.694); interaction effects were not found (F_[1, 21]_ = 2.4, p = 0.136, η²p = 0.102). *Post-hoc* tests found that AE-PG increased activity in both stressed and unstressed animals (t_[df = 21]_ = -6.903, p < 0.001, *d* = -2.788)(Figure 8B).

**Figure 8.**
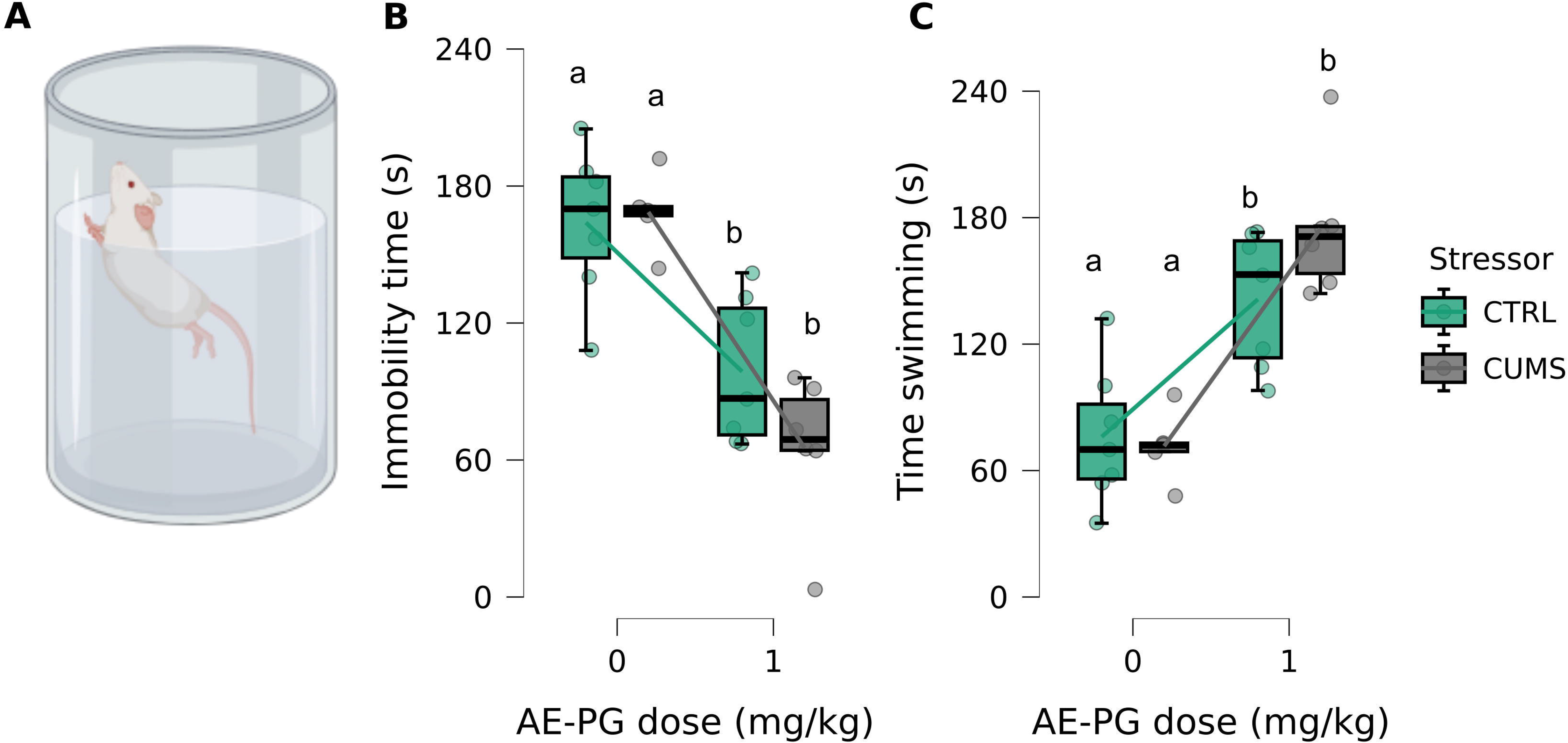
Behavioral effects of AE-PG (1 mg/kg) on mice (*Balb-c*) previously exposed to chronic unpredictable mild stress (CUMS) and tested in the forced swim test (FST). (A) Schematics of the test apparatus. (B) Time spent immobile. (C) Time spent swimming. Letters above each boxplot represent statistically significant differences at the p < 0.05 level; different letters represent significant differences, while groups with equal letters do not differ at the p < 0.05 level. Boxplots represent median ± interquartile range; the lines between boxes connect group averages.

### 3.2. Network topology

The topological architecture of the network (Figure 9) exhibits a classic scale-free, hub-and-spoke organization, typical of biological interactomes. The degree centrality distribution is heavily skewed, yielding a distinct hierarchical structure (Figure 9). The nodes AKT1, INS, and ALB exhibit the highest degree centrality (deep purple nodes in Figure 9), and represent primary central hubs in the network. AKT1 acts as the structural bottleneck of the network, connecting to multiple signaling axes. These hubs represent potential master regulators of the system. A dense, heavily interconnected core surrounds the primary hubs, representing secondary integrators (teal to green nodes in Figure 9). Nodes such as ESR1, ESR2, BDNF, APP, ACE, IGF1, and IRS1 display intermediate-to-high connectivity (Figure 9). This central cluster bridges discrete cellular signaling cascades. Peripheral effectors (yellow to light green nodes in Figure 9) show nodes with minimal connectivity. Proteins such as TNNT2, AURKB, MAOA, GUSB, and OGA act as network leaves, representing highly specialized downstream effectors that depend entirely on upstream signaling from the core hubs.

**Figure 9.**
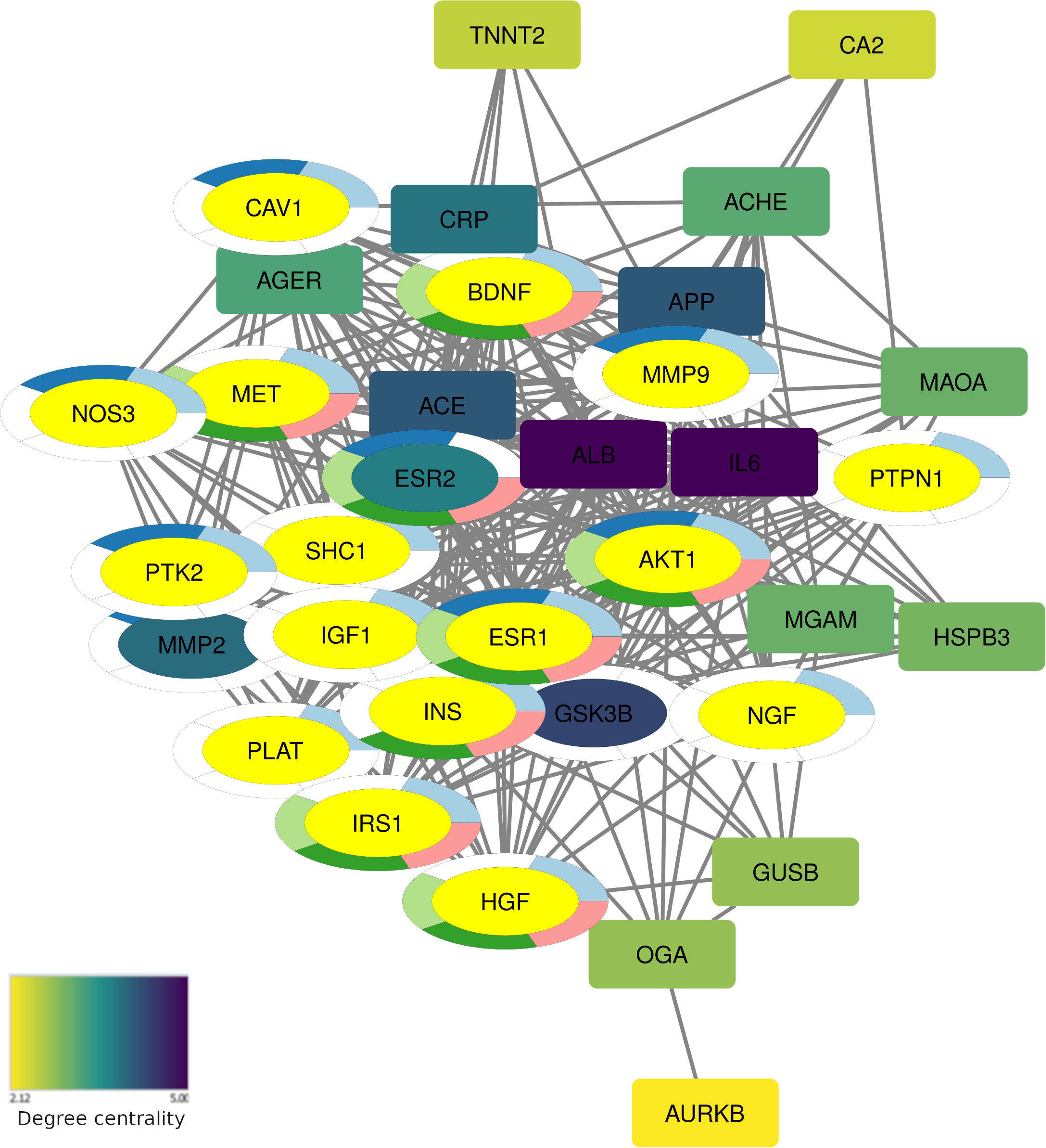
Protein–protein interaction network of potential targets of EA and P3OG associated with anxiety-related pathways. Nodes represent individual proteins/genes, and edges denote predicted or experimentally validated interactions. Node color indicates degree centrality as a continuous variable, ranging from yellow (lowest connectivity) to dark purple (highest connectivity), identifying AKT1, INS, and ALB as the primary topological hubs. Colored oval overlays denote enriched biological functions: light blue (Signaling by Receptor Tyrosine Kinases), dark blue (Extranuclear Estrogen Signaling), light green (PI3K/Akt signaling), dark green (PI5P, PP2A and IER3 Regulate PI3K/AKT Signaling), and pink (Negative regulation of the PI3K/AKT network). The central enrichment of PI3K/Akt signaling, combined with the high connectivity of ESR1 and the primary hubs, highlights the network architecture underlying the stress-dependent homeostatic restabilization model proposed for these phytochemicals.

**Figure 10.**
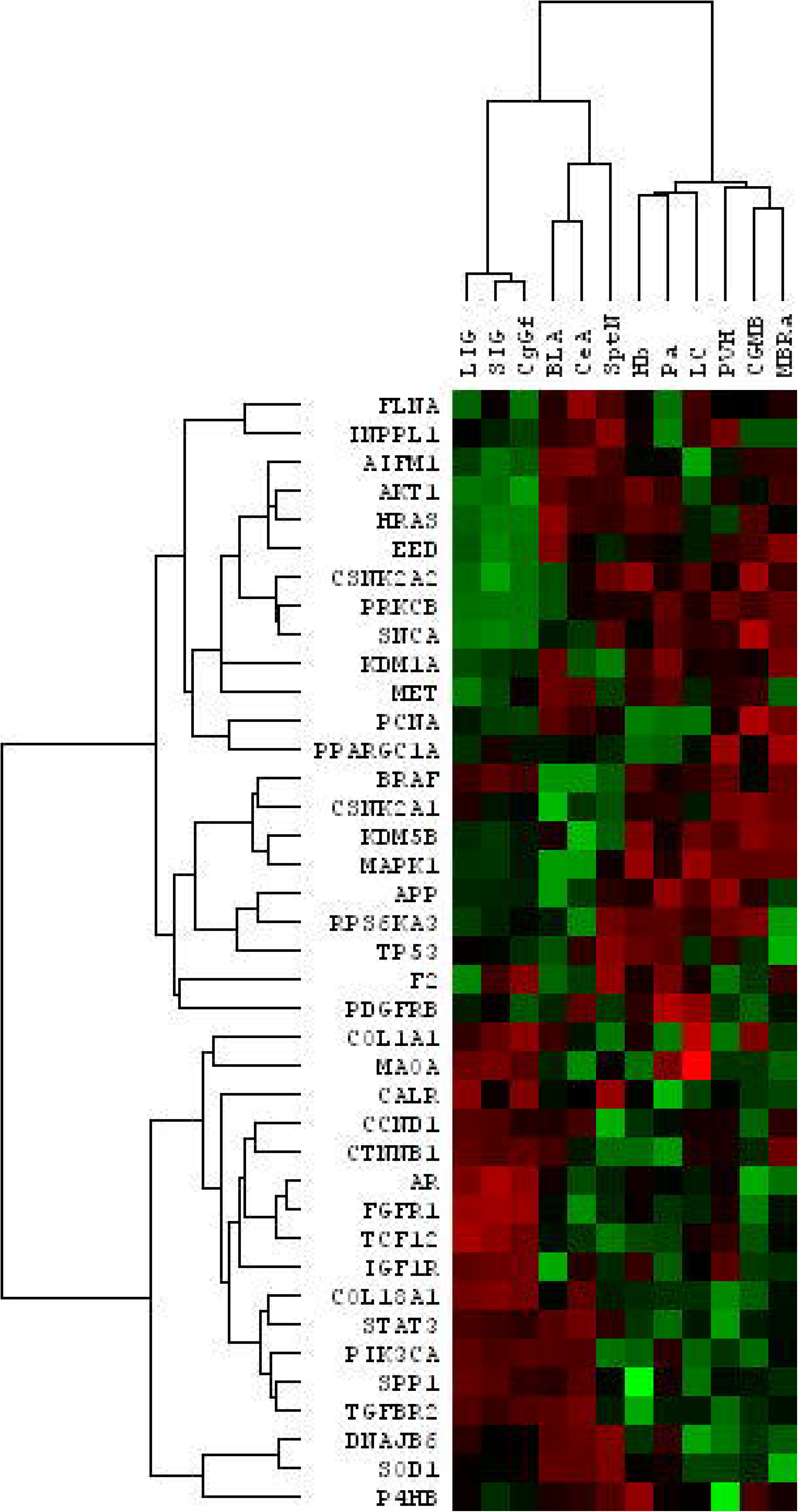
Hierarchical clustering of network-derived genes across anxiety-relevant limbic regions. Regional expression data for the 39 genes comprising the protein–protein interaction network of putative ellagic acid and P3OG targets were retrieved from the Allen Brain Atlas microarray dataset (naïve adult tissue) for 12 limbic-associated regions (LIG, SIG, CgGf, BLA, CeA, SptN, Hb, Pa, PVH, CGMB, MBRa, LC). The per-gene normalized matrix (signed-log transform with gene-centering) was clustered in Cluster 3.0 using a correlation-based similarity with centroid linkage, and the clustered heatmap with both dendrograms was visualized in Java TreeView. Rows are genes and columns are brain regions, both arranged in dendrogram order. The colour scale is red–green, diverging from green (above the gene mean) through black to red (below the gene mean); because normalization was per-gene, the heatmap encodes relative regional patterns within each gene and not absolute expression levels.

**Figure 11.**
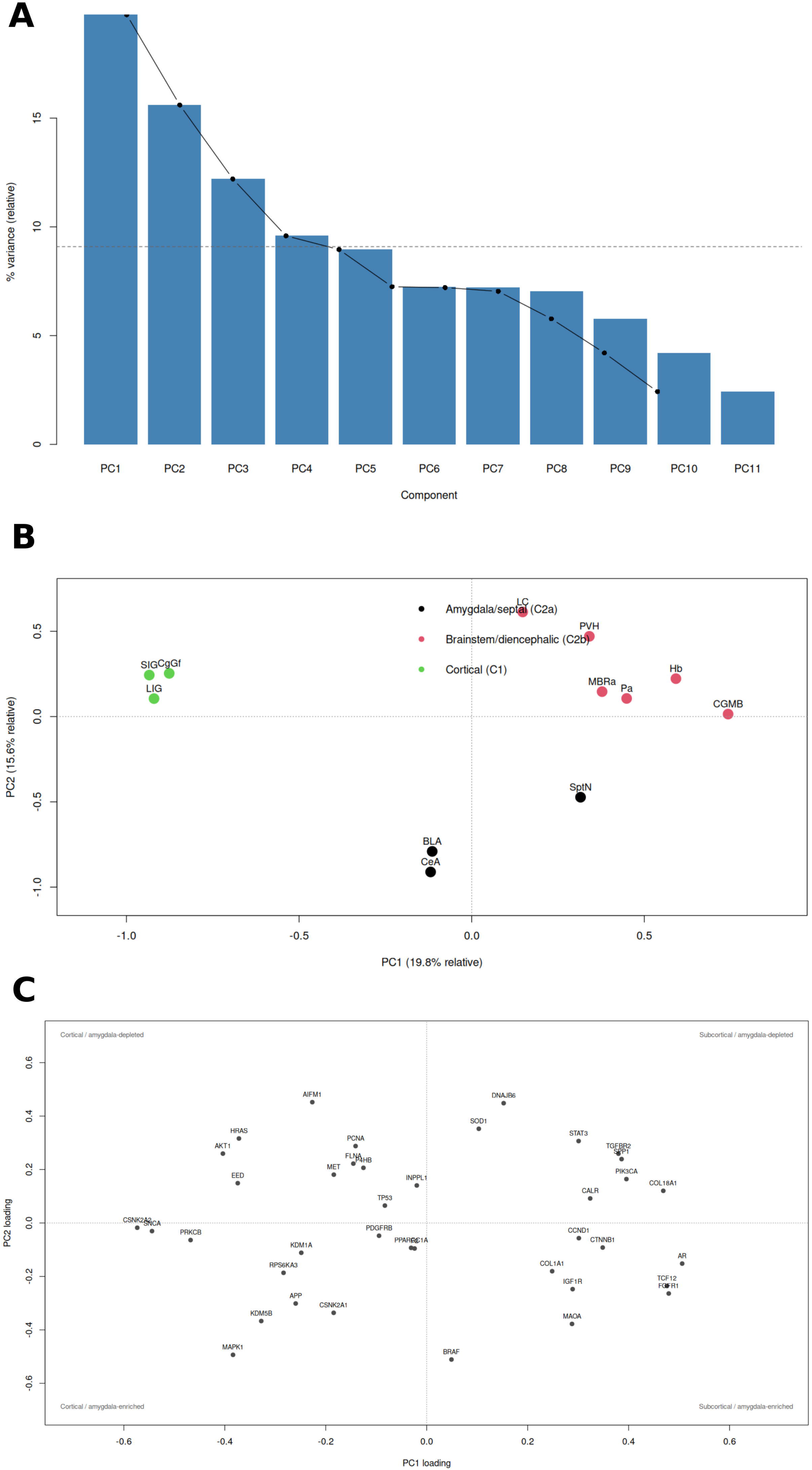
Principal component analysis of network-derived genes across anxiety-relevant limbic regions. Regional expression data for the 39 genes comprising the protein–protein interaction network of putative ellagic acid and P3OG targets were retrieved from the Allen Brain Atlas microarray dataset (naïve adult tissue) for 12 limbic-associated regions (LIG, SIG, CgGf, BLA, CeA, SptN, Hb, Pa, PVH, CGMB, MBRa, LC). The normalized matrix was subjected to PCA in Cluster 3.0. (A) Scree plot of the 11 non-trivial components, showing eigenvalues (bars) and relative percentage of variance (line). The dashed line marks the broken-stick reference; only PC1 (eigenvalue 2.00) and PC2 (1.58) exceed it, together accounting for ∼35% of the relative variance. (B) Score plot of PC1 versus PC2. Each point represents one region, color-coded by hierarchical cluster assignment: cortical (C1: LIG, SIG, CgGf; red), amygdala/septal (C2a: BLA, CeA, SptN; green), and brainstem/diencephalic (C2b: Hb, Pa, PVH, CGMB, MBRa, LC; blue). PC1 separates the cortical regions (scores −0.88 to −0.93) from the brainstem/diencephalic regions (+0.34 to +0.74), with the amygdala and septum intermediate (−0.12 to +0.32); PC2 defines an orthogonal axis on which CeA, BLA, and SptN score negatively (−0.91 to −0.47) and LC and PVH positively (+0.61, +0.47). The spatial segregation of the three color-coded groups reproduces the independently derived hierarchical clustering. (C) Loading plot of PC1 versus PC2 for the same decomposition. Each point represents one gene; quadrant descriptors in grey indicate the regional interpretation of each loading direction (upper left, cortical and amygdala/septal-depleted, e.g. *AKT1*, *HRAS*, *EED*; upper right, subcortical and amygdala/septal-depleted, e.g. *PIK3CA*, *SPP1*, *TGFBR2*; lower left, cortical and amygdala/septal-enriched, e.g. *SNCA*, *CSNK2A2*, *PRKCB*, *MAPK1*; lower right, subcortical and amygdala/septal-enriched, e.g. *AR*, *FGFR1*, *TCF12*, *IGF1R*). Genes near the origin (*INPPL1*, *F2*, *PPARGC1A*, *PDGFRB*, *TP53*) show little regional structure. The tight grouping of *AR*, *FGFR1*, and *TCF12* in the lower-right quadrant corresponds to their co-clustering in the gene dendrogram (*AR*–*FGFR1*, *r* = 0.911). Common to all panels: values are per-gene normalized (signed-log transform with gene-centering), so relative regional patterns are preserved but absolute expression levels are not recoverable; variance percentages are relative to the retained components; and the gene set is network-derived rather than genome-wide, so the two-module structure is conditional on that selection.

### 3.3. Functional enrichment

Gene ontology (GO) enrichment analysis of the dataset reveals a strong functional convergence on the PI3K/Akt signaling axis and related modulatory pathways. Statistical details are found in Table S1. These enriched biological functions are mapped to the network via colored ovals surrounding the gene nodes, identifying the specific pathway membership of each protein:

1. Signaling by Receptor Tyrosine Kinases (Light blue ovals): This pathway characterizes the upstream transduction of growth factor signals. Nodes belonging to this functional group include MET, SHC1, IRS1, IGF1, NOS3, NGF, and PTK2, connecting the extracellular environment to the internal kinase cascades.
2. Extranuclear Estrogen Signaling (Dark blue ovals): Enriched predominantly on ESR1 and ESR2, this functional group illustrates the non-genomic actions of estrogen receptors, which directly interface with intracellular kinases to rapidly alter cellular excitability and survival.
3. PI3K/Akt Signaling (Light green ovals): This is the core canonical pathway represented in the network. The pathway membership prominently includes the central hubs and their immediate modulators: AKT1, GSK3B, INS, and IGF1.
4. PI5P, PP2A and IER3 Regulate PI3K/AKT Signaling (Dark green ovals): Nodes mapped to this auxiliary regulatory pathway (including AKT1 and GSK3B) indicate that the core PI3K/Akt cascade is subject to additional fine-tuning by secondary regulatory proteins (PI5P, PP2A, IER3).
5. Negative Regulation of the PI3K/AKT Network (Pink ovals): Nodes bearing pink ovals, specifically AKT1, INS, and ALB, are implicated in the feedback inhibition mechanisms that dampen PI3K/Akt signaling. This highlights a critical functional balance, preventing unchecked pathway overactivation which could lead to neurotoxicity or pathological plasticity.

### 3.4. Hierarchical clustering of brain areas and genes

The array dendrogram resolved two major groups separated at *r* = −0.971, the strongest split in the data. Group C1 comprised the three cortical regions LIG, SIG, and CgGf (SIG and CgGf merge at *r* = 0.883; LIG joins at *r* = 0.832). Group C2 comprised the remaining nine regions, subdivided into an amygdala/septal cluster (BLA, CeA, SptN; BLA and CeA merge at *r* = 0.474, SptN joins at *r* = 0.105) and a brainstem/diencephalic cluster (Hb, Pa, PVH, CGMB, MBRa, LC, with nested sub-structure: MBRa and CGMB merge at *r* = 0.403; PVH joins at *r* = 0.262; Hb and Pa merge at *r* = 0.301; LC joins at *r* = 0.292). LC was repositioned from its original last column position to sit between Pa and PVH after normalization, indicating that its normalized profile is more similar to Hb and Pa than to the hypothalamic/midbrain block. SptN emerged as a regional outlier carrying a proliferative/extracellular matrix signature (*CCND1*, *CTNNB1*, *COL1A1*, *PIK3CA*, and *TCF12*) that appears nowhere else.

The gene dendrogram resolved two major anti-correlated modules (root merge at *r* = −1.000), indicating that a single dominant axis accounts for the majority of gene-level structure. Module G1 (forebrain-enriched) contained 22 genes and was internally organized into three sub-clusters: G1a1 (*AIFM1*, *AKT1*, *HRAS*, *EED*, *CSNK2A2*, *PRKCB*, *SNCA*; merge correlations 0.79–0.86), G1a2 (*KDM1A*, *MET*, *PCNA*, *PPARGC1A*), and G1b (*BRAF*, *CSNK2A1*, *KDM5B*, *MAPK1*, *APP*, *RPS6KA3*, *TP53*, *F2*, *PDGFRB*). Module G2 (brainstem/subcortical-enriched) contained 17 genes and included a tight extracellular matrix/inflammatory sub-cluster (*PIK3CA*, *SPP1*, *TGFBR2*, *COL18A1*, *STAT3*; merge correlations 0.74–0.84). The single tightest gene pair in the entire tree was *AR*–*FGFR1* (*r* = 0.911). Notably, the PI3K/Akt node genes did not co-cluster: *AKT1* and *HRAS* fell in G1a1, *PIK3CA* in G2, and *INPPL1* with *FLNA* in G1, indicating that these pathway components are not transcriptionally co-regulated across the 12 regions.

To confirm these results, a PCA was run on the same dataset. The first two PCs accounted for only ∼35% of the relative variance (PC1 = 1.999, PC2 = 1.579), and the first five for ∼66% (PC3 = 1.235, PC4 = 0.970, PC5 = 0.907), indicating that the dataset is not low-dimensional. PC1 reproduced the hierarchical forebrain/brainstem split exactly: LIG, SIG, and CgGf scored strongly negative (−0.88 to −0.93); BLA and CeA were near zero (−0.11, −0.12); Hb, Pa, PVH, CGMB, and MBRa scored positive (+0.34 to +0.74). The corresponding gene loadings separated the same G1 (*SNCA*, *CSNK2A2*, *PRKCB*, *AKT1*, *HRAS*, *MAPK1*, *EED*) and G2 (*AR*, *FGFR1*, *TCF12*, *COL18A1*, *PIK3CA*, *SPP1*, *TGFBR2*) modules. PC2 defined an orthogonal amygdala/septal axis, with CeA, BLA, and SptN scoring negatively (−0.91 to −0.47) and LC and PVH positively (+0.61, +0.47); the genes tracking this axis were *BRAF*, *MAPK1*, *MAOA*, *KDM5B*, *CSNK2A1*, *APP*, *FGFR1*, and *IGF1R*, while *AKT1*, *HRAS*, *AIFM1*, and *DNAJB6* were amygdala-depleted. PC3 isolated SptN (opposed to MBRa), driven by *PPARGC1A*, *PCNA*, *CTNNB1*, and *CCND1*. The array-PCA and gene-PCA outputs agreed to within ∼1% on eigenvalues, confirming that the two decompositions describe the same structure.

### 3.4. Molecular docking

Among the five best models of molecular interaction, docking of EA on the binding domain of ERβ produced an interaction with a calculated affinity of -8.556 kcal/mol (Figure 12A). This best-scoring EA pose places the ligand in the pocket such that two hydroxyl groups at the C8 position donate hydrogen bonds to the imidazole side chain of HIS475, while the carbon skeleton engages in hydrophobic contacts with PHE356 and LEU476.

**Figure 12.**
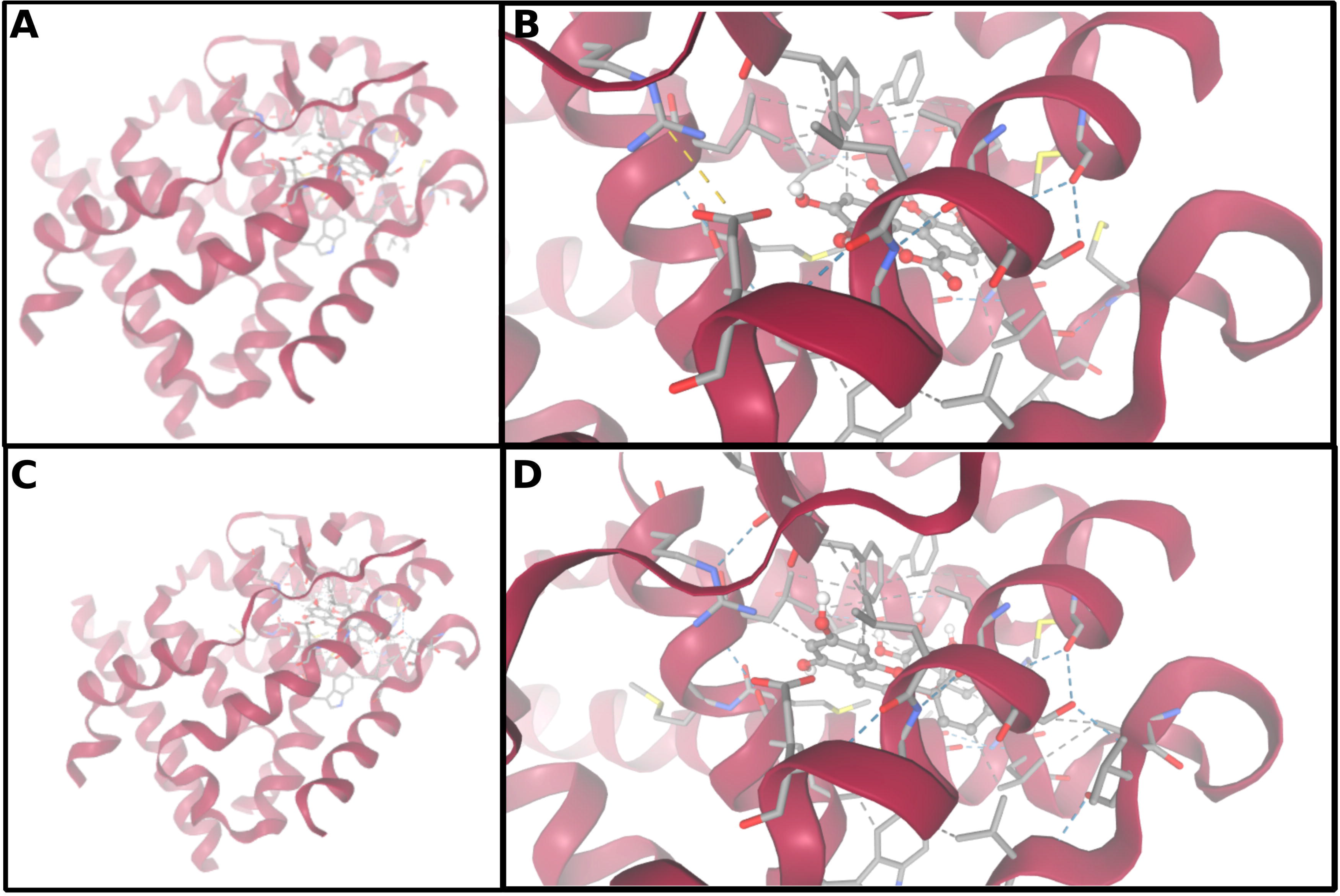
Molecular docking of ellagic acid and pelargonidin-3-O-glucoside into the target binding site. Docked poses are shown for ellagic acid (EA; panels A and B) and pelargonidin-3-O-glucoside (P3OG; panels C and D). Panels A and C show the ligand within the binding cavity in the context of the surrounding protein surface, whereas panels B and D are magnified views of the binding site highlighting the individual ligand–residue interactions. Ligands are rendered as sticks (carbon, grey; oxygen, red; hydrogen, white), and interacting residues are shown as sticks; hydrogen bonds are indicated by dashed lines and hydrophobic contacts by the proximity of non-polar ligand carbons to the side chains of the listed residues.

Three models converged for the interaction between P3OG and the ERβ, binding domain all with low affinity (from 0.73 to -2.169 kcal/mol). The best-scoring P3OG pose, with a calculated binding affinity of −2.169 kcal/mol (Figure 12C), retains a single hydrogen bond between one C8 hydroxyl group and HIS475, and establishes hydrophobic contacts between its carbon atoms and LEU298, LEU301, LEU476, and PHE356. The two ligands therefore share the same anchoring histidine (HIS475) and two of the same hydrophobic partners (PHE356, LEU476), but EA forms a bidentate hydrogen-bond network at this residue whereas P3OG forms only a monodentate interaction; P3OG makes two additional hydrophobic contacts (LEU298, LEU301) that do not compensate for the loss of the second hydrogen bond, consistent with its ∼6.4 kcal·mol⁻¹ lower predicted affinity. Docking scores are computational estimates and require experimental validation; the poses shown are the top-ranked solutions from the docking run and do not by themselves establish binding *in vivo*.

## 4. Discussion

### 4.1. Cross-Species Stress-Dependent Effects of AE-PG

The present work tested the effects of a single acute dose of pomegranate aqueous extract (AE-PG) on anxiety-like behavior in two fish species (lambari and zebrafish) that were subjected to acute restraint stress (ARS) and both anxiety-like and depressive-like behavior in mice that were subjected to unpredictable chronic stress (CUMS). The anxiolytic-like profile of AE-PG was largely stress-dependent and assay-specific. In teleosts, acute restraint stress induced clear anxiogenic-like behaviors in both lambaris and zebrafish, which AE-PG prevented in the novel tank test (normalizing geotaxis, swimming speed, and erratic swimming) and the light/dark test. However, in non-stressed fish, AE-PG produced no measurable behavioral alterations. This pattern aligns with previous reports indicating that pomegranate extracts predominantly reverse anxiety-like behaviors in the context of pre-existing physiological or environmental stressors, such as lead neurotoxicity (Gadouche et al., 2020), cerebral ischemia (Rafieirad & Abbaszadeh, 2017), or metabolic dysfunction (Estrada-Camarena et al., 2020), but remain inert in homeostatic conditions (Flores-Bazán et al., 2023; Estrada-Camarena et al., 2024). The convergence of these findings suggests that the extract’s anxiolytic action may rely on its abilty to mitigate oxidative stress and restore central nervous system balance specifically when it has been disrupted by an allostatic load.

In mice exposed to chronic unpredictable mild stress (CUMS), the stress-dependency of AE-PG’s anxiolytic properties persisted, though its efficacy varied across behavioral assays. The extract successfully reversed CUMS-induced reductions in open arm exploration in the elevated plus-maze (EPM). Conversely, in the open field test, CUMS produced a marked anxiogenic-like profile characterized by increased thigmotaxis, decreased locomotion, reduced stereotypy, and decreased rearing; these deficits were not ameliorated by AE-PG treatment. This distinction indicates that the anxiolytic actions of AE-PG do not represent a generalized dampening of anxiety-related behaviors. Instead, they appear to act on specific stress-sensitive circuits that are engaged by certain test paradigms (such as the EPM and the novel tank test) while sparing others (like thigmotaxis and general locomotion in the OFT).

In contrast to its context-dependent anxiolytic effects, AE-PG exhibited a broader and more consistent antidepressant-like profile. In the forced swim test (FST), the extract significantly reduced behavioral despair in both CUMS-stressed and non-stressed mice, demonstrating baseline efficacy not gated by prior chronic stress exposure; however, the FST is a stronger stressor per se. These observations are consistent with an extensive body of literature showing that various pomegranate preparations (including methanolic, aqueous, and hydroethanolic extracts) reliably reduce immobility in the FST and tail suspension test in rodents, often with efficacy comparable to standard antidepressants like fluoxetine and imipramine (Cervantes-Anaya et al., 2022, 2025; Rahman et al., 2015; Vega-Rivera et al., 2023). This dichotomy suggests that AE-PG’s antidepressant mechanisms are fundamentally distinct from, or at least broader than, those governing its stress-gated anxiolytic actions. Consistent with this, in the splash test, indicative of apathy-like behavior, unstressed mice did not respond to AE-PG treatment.

Mechanistically, previous studies in ovariectomized female rats have demonstrated that AE-PG’s antidepressant-like effects are mediated through estrogen receptor β (ERβ) activation and modulation of the serotonergic system, actions which are blocked by the ER antagonist tamoxifen (Valdés-Sustaita et al., 2017, 2021). Given that the present behavioral assessments used exclusively female mice, and the sex of the fish was undetermined, the observed effects in this study cannot be entirely attributed to phytoestrogenic ERβ signaling. This implies that the extract’s bioactive compounds, particularly ellagic acid and punicalagin, exert polypharmacological effects through alternative pathways. These likely include direct engagement of GABA_A_ receptors, activation of PPARγ (which has been linked to both metabolic and anxiolytic properties), or direct antioxidant and anti-inflammatory actions acting independently of sex hormone receptors (Estrada-Camarena et al., 2024). The persistence of both antidepressant and stress-gated anxiolytic effects across sexes suggests that AE-PG enhances stress resilience via multiple convergent cellular targets, though the precise relative contributions of these pathways in male subjects warrant further investigation.

### 4.2. Integrating Topology, Function, and Stress-Dependent Efficacy

A network pharmacology approach revealed a network of protein interactions that likely represent regulatory mechanisms in response to EA and P3OG, the most likely candidate molecules for the psychoactive effects of AE-PG. Notably, previous studies have demonstrated that ellagic acid (EA), the principal bioactive of *Punica granatum*, can directly modulate the PI3K/Akt signaling axis in various cellular stress models (e.g., Aldayel et al., 2025). The topological and functional architecture of this network provides a compelling mechanistic framework for understanding why the anxiolytic actions of EA and P3OG are strictly stress-dependent. The behavioral evidence from zebrafish, lambari, and mice demonstrates that these compounds exhibit no benefit in unstressed animals, yet significantly reduce anxiety-related behaviors in stressed subjects.

The enrichment of Extranuclear Estrogen Signaling within the dense network core expands on the earlier pharmacological implication of estrogen receptors ESR1 and ESR2 (Valdés-Sustaita et al., 2021). The positioning of ESR1 allows it to act as an upstream rheostat that rapidly engages the PI3K/Akt cascade via crosstalk with AKT1 and GSK3B. This interaction is critical in the stressed brain, where excessive glucocorticoid signaling dysregulates normal synaptic function (Popoli et al., 2011). The co-enrichment of both positive PI3K/Akt signaling and negative PI3K/Akt regulation reveals a tightly balanced homeostatic control mechanism. In unstressed animals, this physiological balance is intact. The canonical pathways (RTK, PI3K/Akt) operate at baseline (Guo et al., 2024), and the negative feedback loops are adequately preventing overexcitation. Because the system is already in equilibrium, EA and P3OG’s subtle modulation of estrogen receptors (Estrada-Camarena et al., 2024; Valdés-Sustaita et al., 2021) lacks a measurable net effect, explaining the lack of behavioral efficacy in non-stressed subjects.

The docking results reveal a striking difference in the predicted binding free energies, with EA (−8.56 kcal/mol) exhibiting a ∼6.4 kcal/mol advantage over P3OG (−2.17 kcal/mol) in simulated interactions with the binding site of ERβ/ESR2. Both ligands engage the same histidine residue and share hydrophobic contacts with PHE and LEU side chains; P3OG even establishes two additional hydrophobic contacts with distinct LEU residues. The main divergence lies in the hydrogen-bonding pattern: EA donates two hydrogen bonds from its hydroxyl groups to the histidine, whereas P3OG forms only a single such interaction. Because a neutral, geometrically optimal hydrogen bond in a protein binding site typically contributes 0.5–1.5 kcal/mol (Bissantz et al., 2010), losing one hydrogen bond alone cannot quantitatively account for the entire affinity gap. However, the bidentate H-bond network of EA may cooperatively pre-organize the ligand and restrict residual mobility within the pocket, thereby yielding a more favorable entropic and enthalpic signature than the sum of its individual contacts (D. Chen et al., 2016). These results align with the network-based analysis, further indicating the role of estrogen signaling and convergence on PI3K/Akt as putative mechanisms of *P. granatum* anxiolytic-like effects.

The clustering and PCA converge on three robust features of the baseline regional expression of the network genes. First, the dominant axis separates a cortical compartment (insula and anterior cingulate) from a brainstem/diencephalic compartment (habenula, paraventricular thalamus, periventricular hypothalamus, central grey, midbrain raphe, and locus coeruleus), with the amygdala and septum occupying an intermediate position. Both methods recover this axis with near-identical boundaries and gene membership, which is the strongest internal validation available for a matrix of this size. Second, the amygdala/septal compartment defines a distinct, orthogonal axis (PC2) enriched for *BRAF*, *MAPK1*, *MAOA*, *KDM5B*, and *FGFR1* and relatively depleted for *AKT1* and *HRAS*. Third, the septal region (SptN) carries a region-specific proliferative/ECM program orthogonal to both major axes. The central implication is that the network genes are transcribed in the anxiety-relevant limbic circuitry examined here. This is a necessary, though not sufficient, condition for the network to be functionally engaged in these regions, and it supports the anatomical plausibility of the EA/P3OG target set.

However, the network genes are not expressed as a single coordinated unit: they partition into two anti-correlated modules, and the PI3K/Akt node itself is transcriptionally split, with *AKT1* and *HRAS* in the cortical module, *PIK3CA* in the subcortical module, and *INPPL1* showing no strong regional structure. This dissociation is informative, as it suggests that the functional convergence on PI3K/Akt identified in the network pharmacology analysis is unlikely to be reflected at the level of mRNA co-regulation, and is more consistent with convergence at the level of protein–protein interaction, post-translational modification (e.g., Akt phosphorylation, PIP_3_ turnover, INPPL1-mediated lipid dephosphorylation), and membrane-proximal complex assembly, all modes of action canonically attributed to extranuclear estrogen signalling (Levin, 2005). In this sense, the regional expression architecture complements the network result: the network describes a signalling-level convergence, and the transcriptome describes a regionally heterogeneous distribution of the participating components.

The tight co-clustering of *AR* with *FGFR1* and their joint placement in the subcortical/amygdala-enriched quadrant of PC1–PC2 space identify a specific candidate substrate for the Extranuclear Estrogen Signaling arm of the network. AR and the estrogen receptors share co-regulators and can assemble into membrane-associated signalling complexes with growth-factor receptors (Lonard & O’Malley, 2014); the co-expression of *AR*, *FGFR1*, and *IGF1R* in the compartment that is transcriptionally distinct from the cortex provides a regional basis for the non-genomic signalling proposed by the network analysis. Interestingly, in zebrafish mutations of the *fgfr1a* result in increased aggression, decreased anxiety, and increased exploration through mechanisms that are mediated by histaminergic neurotransmission (Norton et al., 2011). Because the behavioural effect of the PG extract is contingent on prior stress, and because the amygdala/septal axis (PC2) dissociates BLA from CeA, the two nuclei most directly implicated in stress-dependent fear versus anxiety output (LeDoux, 2000, 2016), the amygdala-enriched PC2 genes (*MAPK1*, *BRAF*, *MAOA*, *FGFR1*) constitute the most plausible regional correlates of the stress-dependency. The docking results add a further constraint at the ligand level, predicting that EA has a higher binding affinity than P3OG. Combined with the regional expression architecture, this predicts that EA is the more likely contributor to the stress-dependent anxiolytic-like effect of the PG extract, and that follow-up experiments should prioritize regional readouts of PI3K/Akt pathway activity rather than *AKT1* or *PIK3CA* transcript levels.

In stressed animals, chronic psychological and physiological strain induces a pathological allostatic load (McEwen, 2007). This can be characterized by the dysregulation of the central hubs (AKT1, INS, ALB), aberrant RTK signaling, and a breakdown of homeostatic negative feedback (Guo et al., 2024). The stress-induced network state becomes biased toward neuroinflammation and excitotoxicity. In this vulnerable state, EA and P3OG leverage the network’s highly connected central hubs. By activating the estrogen receptor axis, the compounds rebuild topological stability. They functionally bias the enriched PI3K/Akt cascade toward cellular resilience while simultaneously re-engaging the homeostatic negative regulatory loops, effectively restoring the network to its pre-stress homeostatic state (Duman & Monteggia, 2006).

The EA/P3OG-anxiety network is governed by a central PI3K/Akt signaling axis that is intimately linked to growth factor signaling (RTKs) and neuroendocrine modulation (ESR1). The topological centrality of AKT1, combined with the functional enrichment of both positive and negative regulatory pathways, provides a mechanistic explanation for the stress-selective efficacy of the plant-derived molecules. Rather than acting as unconditional neuromodulators, EA and P3OG function as homeostatic re-stabilizers, responding to and correcting the pathological shifts induced by stress, while remaining inert within balanced, unstressed biological systems.

## 5. Conclusions and Limitations

This work has limitations that should be considerer when interpreting the results. The mouse experiments included only female mice and the sex of the fish was not determined, limiting inferences about sex-specific and phytoestrogenic mechanisms, although in other studies the antidepressant-like effects of the AE-PG in male mice have been reported (Cervantes-Anaya et al 2025). The behavioral conclusions are based on selected assays and acute extract administration in fish; dose-response, chronic treatment, and additional behavioral domains were not examined. The proposed mechanisms rely on network pharmacology and molecular docking and therefore require direct *in vivo* validation of ERβ/PI3K/Akt signaling, antioxidant/anti-inflammatory actions, and the specific contributions of ellagic acid and punicalagin; therefore, they should be interpreted as hypothesis to be tested empirically. Finally, the use of different stress models across species and the absence of AE-PG effects on some open-field parameters caution against broad generalizations. In addition, in mice, a behavioral battery was used to meet the 3R principle, and cumulative effects of one test on another could not be ruled out; however, the order followed for each test was taken into account in relation to the stress level (Sánchez-Cortes et al 2026).

Overall, AE-PG produced anxiolytic-like effects that were largely stress-dependent and assay-specific: it prevented acute restraint stress-induced anxiety-like behavior in zebrafish and lambari and reversed CUMS-induced open-arm avoidance in mice, but it did not alter behavior in unstressed fish or ameliorate all CUMS-induced open-field deficits. In contrast, AE-PG showed a broader antidepressant-like profile in the FST in both stressed and non-stressed mice. These findings suggest that AE-PG acts as a “homeostatic restabilizer” rather than an unconditional neuromodulator, with network pharmacology and docking implicating estrogen receptor signaling and the PI3K/Akt pathway as convergent mechanisms through which ellagic acid and punicalagin may restore stress-disrupted networks.

## References

Abdel Moneim, A. E. (2012). Evaluating the Potential Role of Pomegranate Peel in Aluminum-Induced Oxidative Stress and Histopathological Alterations in Brain of Female Rats. Biological Trace Element Research, 150(1), 328–336. 10.1007/s12011-012-9498-2

Abu-Taweel, G. M., & Al-Mutary, M. G. (2021). Pomegranate juice moderates anxiety- and depression-like behaviors in AlCl_3_-treated male mice. Journal of Trace Elements in Medicine and Biology, 68, 126842. 10.1016/j.jtemb.2021.126842

Aldayel, T. S., Haleka, S. A. A., Ebaid, H. M., EL-Fadeal, N. M. A., Abdelrazek, H. M., Gad EL-Hak, H. N., Darwish, K. M., & Rashwan, H. M. (2025). Ellagic acid mediates cardioprotection against adrenaline-induced toxicity via PI3K/AKT and Keap1-NRF2 axes. Scientific Reports, 15(1), 36489. 10.1038/s41598-025-21317-7

Aleksandrova, S., Alexova, R., Dragomanova, S., Kalfin, R., Nicoletti, F., Fagone, P., Petralia, M. C., Mangano, K., & Tancheva, L. (2023). Preventive and Therapeutic Effects of Punica granatum L. Polyphenols in Neurological Conditions. International Journal of Molecular Sciences, 24(3), 1856. 10.3390/ijms24031856

Alp, D. (2025, agosto 29). The Pomegranate in History and Myth. JSTOR Daily. https://daily.jstor.org/the-pomegranate-in-history-and-myth/

Bardou, P., Mariette, J., Escudié, F., Djemiel, C., & Klopp, C. (2014). jvenn: An interactive Venn diagram viewer. BMC Bioinformatics, 15(1), 293. 10.1186/1471-2105-15-293 [Website: https://www.bioinformatics.com.cn/static/others/jvenn/example.html]

Bencan, Z., Sledge, D., & Levin, E. D. (2009). Buspirone, chlordiazepoxide and diazepam effects in a zebrafish model of anxiety. Pharmacology Biochemistry & Behavior, 94(1), 75–80. 10.1016/j.pbb.2009.07.009

Bissantz, C., Kuhn, B., & Stahl, M. (2010). A Medicinal Chemist’s Guide to Molecular Interactions. Journal of Medicinal Chemistry, 53(14), 5061–5084. 10.1021/jm100112j

Bugnon, M., Röhrig, U. F., Goullieux, M., Perez, M. A. S., Daina, A., Michielin, O., & Zoete, V. (2024). SwissDock 2024: Major enhancements for small-molecule docking with Attracting Cavities and AutoDock Vina. Nucleic Acids Research, 52(W1), W324–W332. 10.1093/nar/gkae300

Cachat, J., Stewart, A., Grossman, L., Gaikwad, S., Kadri, F., Chung, K. M., Wu, N., Wong, K., Roy, S., Suciu, C., Goodspeed, J., Elegante, M., Bartels, B., Elkhayat, S., Tien, D., Tan, J., Denmark, A., Gilder, T., Kyzar, E., … Kalueff, A. V. (2010). Measuring behavioral and endocrine responses to novelty stress in adult zebrafish. Nature Protocols, 5(11), 1786–1799. 10.1038/nprot.2010.140

Cervantes-Anaya, N., Aedo-Torrado, A., Estrada-Camarena, E., Cruz, V. P. de la, Ortega, D. R., González-Trujano, M. E., & López-Rubalcava, C. (2025). Role of Nitric Oxide in the Antidepressant Effect of an Aqueous Extract of Punica granatum L.: Effects on GSH/GSSG Ratio and Lipoperoxidation in Adult Male Swiss Webster Mice. International Journal of Molecular Sciences, 26(21). 10.3390/ijms262110255

Cervantes-Anaya, N., Azpilcueta-Morales, G., Estrada-Camarena, E., Ramírez Ortega, D., Pérez de la Cruz, V., González-Trujano, M. E., & López-Rubalcava, C. (2022). Pomegranate and Its Components, Punicalagin and Ellagic Acid, Promote Antidepressant, Antioxidant, and Free Radical-Scavenging Activity in Ovariectomized Rats. Frontiers in Behavioral Neuroscience, 16, 836681. 10.3389/fnbeh.2022.836681

Chen, D., Oezguen, N., Urvil, P., Ferguson, C., Dann, S. M., & Savidge, T. C. (2016). Regulation of protein-ligand binding affinity by hydrogen bond pairing. Science Advances, 2(3), e1501240. 10.1126/sciadv.1501240

Chen, P., Guo, Z., Lei, J., & Wang, Y. (2024). Pomegranate polyphenol punicalin ameliorates lipopolysaccharide-induced memory impairment, behavioral disorders, oxidative stress, and neuroinflammation via inhibition of TLR4-NF-кB pathway. Phytotherapy Research, 38(7), 3489–3508. 10.1002/ptr.8219

Conselho Nacional de Controle de Experimentação Animal - CONCEA. (2024). Diretriz brasileira para o cuidado e a utilização de animais em atividades de ensino ou de pesquisa científica—DBCA (M. dos S. Gonçalves & A. Am. B. Viana, Org.). Ministério da Ciência, Tecnologia e Inovação. https://ceua.ufes.br/sites/ceua.ufes.br/files/field/anexo/dbca_2024_0.pdf

de Oliveira, C. M., Gressler, L. T., Ferreira, M. V. dos A. L., & Whitton, R. G. M. (2023). Capítulo 5—Peixes I: Lambari, tilápia e zebrafish. Em L. M. F. Winter, *Guia brasileiro de produção, manutenção ou utilização de animais em atividades de ensino ou pesquisa científica/Concea* (p. 330–383). Ministério da Ciência, Tecnologia e Inovação.

De Sousa, B. L., Chaves, S. N., Albuquerque, E., Rodrigues, J., Coimbra, V., Miranda, S., Caldas, A. L., Leite, M., Dos Santos, M. P., Côrrea Filho, R. A. C., Santos, A. D. F. D., Maximino, C., & Siqueira-Silva, D. (2024). Gametogenesis and seminatural reproduction of the Amazon twospot astyanax Astyanax bimaculatus (Linnaeus, 1758) cultivated in an enriched environment. Animal Reproduction Science, 267, 107522. 10.1016/j.anireprosci.2024.107522

Devi, T., & Das, S. (2016). Evaluation of the protective effect of ethanolic extract of leaves of Punica granatum Linn. On forced swimming induced chronic fatigue syndrome in mice. International Journal of Pharmacy and Pharmaceutical Sciences, 9, 207. 10.22159/ijpps.2017v9i1.13712

Dhingra, D., & Chhillar, R. (2012). Antidepressant-like activity of ellagic acid in unstressed and acute immobilization-induced stressed mice. Pharmacological Reports, 64(4), 796–807. 10.1016/S1734-1140(12)70875-7

Duman, R. S., & Monteggia, L. M. (2006). A neurotrophic model for stress-related mood disorders. Biological Psychiatry, 59(12), 1116–1127. 10.1016/j.biopsych.2006.02.013

Eberhardt, J., Santos-Martins, D., Tillack, A. F., & Forli, S. (2021). AutoDock Vina 1.2.0: New Docking Methods, Expanded Force Field, and Python Bindings. Journal of Chemical Information and Modeling, 61, 3891–3898. 10.1021/acs.jcim.1c00203

Egan, R. J., Bergner, C. L., Hart, P. C., Cachat, J. M., Canavello, P. R., Elegante, M. F., Elkhayat, S. I., Bartels, B. K., Tien, A. K., Tien, D. H., Mohnot, S., Beeson, E., Glasgow, E., Amri, H., Zukowska, Z., & Kalueff, A. V. (2009). Understanding behavioral and physiological phenotypes of stress and anxiety in zebrafish. Behavioural Brain Research, 205(1), 38–44. 10.1016/j.bbr.2009.06.022

Estrada-Camarena, E. M., López-Rubalcava, C., Ramírez-Rodríguez, G. B., Pulido, D., Cervantes-Anaya, N., Azpilcueta-Morales, G., Granados-Juárez, A., Vega-Rivera, N. M., Islas-Preciado, D., Treviño, S., de Gortari, P., González-Trujano, M. E., & García-Viguera, C. (2020). Aqueous extract of pomegranate enriched in ellagitannins prevents anxiety-like behavior and metabolic changes induced by cafeteria diet in an animal model of menopause. Neurochemistry International, 141, 104876. 10.1016/j.neuint.2020.104876

Estrada-Camarena, E. M., López-Rubalcava, C., Vega-Rivera, N. M., & González-Trujano, M. E. (2024). Antidepressant- and Anxiolytic-like Effects of Pomegranate: Is It Acting by Common or Well-Known Mechanisms of Action? Plants, 13(16). 10.3390/plants13162205

Flores-Bazán, T., Betanzos-Cabrera, G., Guerrero-Solano, J. A., Negrete-Díaz, J. V., German-Ponciano, L. J., & Olivo-Ramírez, D. (2023). Pomegranate (*Punica granatum* L.) and its phytochemicals as anxiolytic; an underreported effect with therapeutic potential: A systematic review. Brain Research, 1820, 148554. 10.1016/j.brainres.2023.148554

Flores-Gutiérrez, E., Cabrera-Muñoz, E. A., Vega-Rivera, N. M., Ortiz-López, L., & Ramírez-Rodríguez, G. B. (2018). Exposure to Patterned Auditory Stimuli during Acute Stress Prevents Despair-Like Behavior in Adult Mice That Were Previously Housed in an Enriched Environment in Combination with Auditory Stimuli. Neural Plasticity, 2018, 8205245. 10.1155/2018/8205245.

Gadouche, L., Djebli, N., & Zerrouki, K. (2020). Algerian pomegranate peel decreases lead concentration in brain and improves neurological disorders. Polish Journal of Natural Sciences, 35, 97–107.

Ghisleni, G., Capiotti, K. M., Silva, R. S., Oses, J. P., Piato, Â. L., Soares, V., Bogo, M. R., & Bonan, C. D. (2012). The role of CRH in behavioral responses to acute restraint stress in zebrafish. Progress in Neuro-Psychopharmacology and Biological Psychiatry, 36, 176–182. 10.1016/j.pnpbp.2011.08.016

González-Trujano, M. E., Pellicer, F., Mena, P., Moreno, D. A., & García-Viguera, C. (2015). Antinociceptive and anti-inflammatory activities of a pomegranate ( *Punica granatum* L.) extract rich in ellagitannins. International Journal of Food Sciences and Nutrition, 66(4), 395–399. 10.3109/09637486.2015.1024208

Guo, N., Wang, X., Xu, M., Bai, J., Yu, H., & Le Zhang. (2024). PI3K/AKT signaling pathway: Molecular mechanisms and therapeutic potential in depression. Pharmacological Research, 206, 107300. 10.1016/j.phrs.2024.107300

Huhn, M., Tardy, M., Spineli, L. M., Kissling, W., Förstl, H., Pitschel-Walz, G., Leucht, C., Samara, M., Dold, M., Davis, J. M., & Leucht, S. (2014). Efficacy of Pharmacotherapy and Psychotherapy for Adult Psychiatric Disorders: A Systematic Overview of Meta-analyses. JAMA Psychiatry, 71(6), 706–715. 10.1001/jamapsychiatry.2014.112

Islam, M. S., Hossain, R., Ahmed, T., Rahaman, M. M., Al-Khafaji, K., Khan, R. A., Sarkar, C., Bappi, M. H., Andrade, E. M. de, Araújo, I. M., Coutinho, H. D. M., Kowalska, G., Kowalski, R., Hanif, M. A., & Islam, M. T. (2022). Anxiolytic-like Effect of Quercetin Possibly through GABA Receptor Interaction Pathway: *In Vivo* and *In Silico* Studies. Molecules, 27(21). 10.3390/molecules27217149

Islas-Preciado, D., López-Rubalcava, C., Estrada-Camarena, E., de Gortari, P., & Castro-García, M. (2023). Effect of chronic unpredictable stress in female Wistar-Kyoto rats subjected to progesterone withdrawal: Relevance for Premenstrual Dysphoric Disorder neurobiology. Psychoneuroendocrinology, 155, 106331. 10.1016/j.psyneuen.2023.106331

Jahromy, M. H., Shariatifar, A., Samiee, S., Vaziri, M., Shahraki, M. B., & Dara, S. M. (2014). Involvement of Serotonergic System and Magnesium on Anxiolytic Effects of Pomegranate in Male Mice. World Journal of Neuroscience, 04(04), 293–298. 10.4236/wjns.2014.44032

Kalueff, A. V., Aldridge, J. W., LaPorte, J. L., Murphy, D. L., & Tuohimaa, P. (2007). Analyzing grooming microstructure in neurobehavioral experiments. Nature Protocols, 2(10), 2538–2544. 10.1038/nprot.2007.367

Kaushal, V., Chand, P., Rupanagunta, G. P., Joshi, R., Nandave, M., & Upadhyay, J. (2024). Investigating the Therapeutic Potential of Medicinal Plants in Managing Mental Health Disorders. Em M. Nandave, R. Joshi, & J. Upadhyay (Org.), Ethnopharmacology and OMICS Advances in Medicinal Plants Volume 2: Revealing the Secrets of Medicinal Plants (p. 515–530). Springer Nature. 10.1007/978-981-97-4292-9_23

Kopcalic, K., Arcaro, J., Pinto, A., Ali, S., Barbui, C., Curatoli, C., Martin, J., & Guaiana, G. (2025). Antidepressants versus placebo for generalised anxiety disorder (GAD). Cochrane Database of Systematic Reviews, (1). 10.1002/14651858.CD012942.pub2

Kysil, E. V., Meshalkina, D. A., Frick, E. E., Echevarria, D. J., Rosemberg, D. B., Maximino, C., Lima, M. G., Abreu, M. S., Giacomini, A. C., Barcellos, L. J. G., Song, C., & Kalueff, A. V. (2017). Comparative Analyses of Zebrafish Anxiety-Like Behavior Using Conflict-Based Novelty Tests. Zebrafish, 14(3), 197–208. 10.1089/zeb.2016.1415

Lawrence, C. (2007). The husbandry of zebrafish (*Danio rerio*): A review. Aquaculture, 269, 1–20. 10.1016/j.aquaculture.2007.04.077

Lee, H. R. (2012). Method for improving pomegranate extract effect for treating anxiety, depression, or concentration reduction (Ministry of Intellectual Property Patent N. KR1020110113018). https://patentscope.wipo.int/search/en/detail.jsf?docId=KR45547107&_cid=P21-MSRGQA-19081-1

Lee, H.-Y. (2011). Pomegranate extract having a high ellagic acid content, and use of the pomegranate extract (US Patent and Trademark Office Patent N. US20130028994). https://patentscope.wipo.int/search/en/detail.jsf?docId=US76681789&_cid=P21-MSRG7P-99692-1

Levin, E. D., Bencan, Z., & Cerutti, D. T. (2007). Anxiolytic effects of nicotine in zebrafish. Physiology & Behavior, 90, 54–58. 10.1016/j.physbeh.2006.08.026

Liu, C., Wang, N., Zhang, Y.-Y., Qu, H., Liu, L.-X., Zhang, L.-H., & Liu, Y.-G. (2025). Pomegranate: Historical origins, nutritional composition, health functions and processing development research. Critical Reviews in Food Science and Nutrition, 65(33), 8447–8469. 10.1080/10408398.2025.2500676

Liu, J., Ning, W., Zhang, N., Zhu, B., & Mao, Y. (2024). Estimation of the Global Disease Burden of Depression and Anxiety between 1990 and 2044: An Analysis of the Global Burden of Disease Study 2019. Healthcare, 12(17), 1721. 10.3390/healthcare12171721

Liu, Q., He, H., Yang, J., Feng, X., Zhao, F., & Lyu, J. (2020). Changes in the global burden of depression from 1990 to 2017: Findings from the Global Burden of Disease study. Journal of Psychiatric Research, 126, 134–140. 10.1016/j.jpsychires.2019.08.002

Loren, D. J., Seeram, N. P., Schulman, R. N., & Holtzman, D. M. (2005). Maternal Dietary Supplementation with Pomegranate Juice Is Neuroprotective in an Animal Model of Neonatal Hypoxic-Ischemic Brain Injury. Pediatric Research, 57(6), 858–864. 10.1203/01.PDR.0000157722.07810.15

Maximino, C., Marques de Brito, T., Dias, C. A. G. de M., Gouveia, A., & Morato, S. (2010). Scototaxis as anxiety-like behavior in fish. Nature Protocols, 5(2), 209–216. 10.1038/nprot.2009.225

Maximino, C., Oliveira, D. L. de, Rosemberg, D. B., Batista, E. de J. O., Herculano, A. M., Oliveira, K. R. M., Benzecry, R., & Blaser, R. (2012). A comparison of the light/dark and novel tank tests in zebrafish. Behaviour, 149(10–12), 1099–1123. 10.1163/1568539X-00003029

McEwen, B. S. (2007). Physiology and neurobiology of stress and adaptation: Central role of the brain. Physiological Reviews, 87(3), 873–904. 10.1152/physrev.00041.2006

Padmanaban, N., Ambosie, R., Choy, S., Marcus, S., Nilsson, S. R. O., Keene, A. C., Kowalko, J. E., & Duboué, E. R. (2025). Automated Behavioral Profiling Using Neural Networks Reveals Differences in Stress-Like Behavior Between Cave and Surface-Dwelling Astyanax mexicanus. Journal of Experimental Zoology Part B: Molecular and Developmental Evolution, 344(6), 352–362. 10.1002/jez.b.23311

Panossian, A. (2017). Understanding adaptogenic activity: Specificity of the pharmacological action of adaptogens and other phytochemicals. Annals of the New York Academy of Sciences, 1401, 49–64. 10.1111/nyas.13399

Parichy, D. M., Elizondo, M. R., Mills, M. G., Gordon, T. N., & Engeszer, R. E. (2009). Normal table of postembryonic zebrafish development: Staging by externally visible anatomy of the living fish. Developmental Dynamics, 238, 2975–3015. 10.1002/dvdy.22113

Pike, A. C. W., Brzozowski, A. M., Hubbard, R. E., Bonn, T., Thorsell, A., Engström, O., Ljunggren, J., Gustafsson, J., & Carlquist, M. (1999). Structure of the ligand-binding domain of oestrogen receptor beta in the presence of a partial agonist and a full antagonist. The EMBO Journal, 18(17), 4608–4618. 10.1093/emboj/18.17.4608

Popoli, M., Yan, Z., McEwen, B., & Sanacora, G. (2011). The stressed synapse: The impact of stress and glucocorticoids on glutamate transmission. Nature reviews. Neuroscience, 13(1), 22–37. 10.1038/nrn3138

Porsolt, R. D., Bertin, A., & Jalfre, M. (1977). Behavioral despair in mice: A primary screening test for antidepressants. Archives Internationales de Pharmacodynamie et de Therapie, 229(2), 327–336.

Rafieirad, M., & Abbaszadeh, H. (2017). Pomegranate seed extract reduces ischemia induced anxiety in male rats. Journal of Herbmed Pharmacology, 6, 85–89.

Rahman, S. A., Kadar, B. A., Baharuldin, M. T. H., & Moklas, M. A. M. (2015). Antidepressant-like effect of methanolic extract of *Punica granatum* (pomegranate) in mice model of depression. Journal of Natural Products and Biomedical Research, 1, 16–20.

Shannon, P., Markiel, A., Ozier, O., Baliga, N. S., Wang, J. T., Ramage, D., Amin, N., Schwikowski, B., & Ideker, T. (2003). Cytoscape: A software environment for integrated models of biomolecular interaction networks. Genome Research, 13(11), 2498–2504. 10.1101/gr.1239303

Steel, Z., Marnane, C., Iranpour, C., Chey, T., Jackson, J. W., Patel, V., & Silove, D. (2014). The global prevalence of common mental disorders: A systematic review and meta-analysis 1980-2013. International Journal of Epidemiology, 43(2), 476–493. 10.1093/ije/dyu038

Syeda, S. F., G., S. P. K., & Mohsin, M. (2020). Evaluations of antidepressant activity of *Punica granatum* peel extract in albino mice. International Journal of Basic & Clinical Pharmacology, 9(3), 449. 10.18203/2319-2003.ijbcp20200720

Valdés-Sustaita, B., Estrada-Camarena, E., González-Trujano, M. E., & López-Rubalcava, C. (2021). Estrogen receptors-β and serotonin mediate the antidepressant-like effect of an aqueous extract of pomegranate in ovariectomized rats. Neurochemistry International, 142, 104904. 10.1016/j.neuint.2020.104904

Valdés-Sustaita, B., López-Rubalcava, C., González-Trujano, M. E., García-Viguera, C., & Estrada-Camarena, E. (2017). Aqueous Extract of Pomegranate Alone or in Combination with Citalopram Produces Antidepressant-Like Effects in an Animal Model of Menopause: Participation of Estrogen Receptors. International Journal of Molecular Sciences, 18(12), Article 12. 10.3390/ijms18122643

Vega-Rivera, N.-M., González-Trujano, M. E., Luna-Angula, A., Sánchez-Chapul, L., & Estrada-Camarena, E. M. (2023). Antidepressant-like effects of the *Punica granatum* and citalopram combination are associated with structural changes in dendritic spines of granule cells in the dentate gyrus of rats. Frontiers in Pharmacology, 14, 1211663. 10.3389/fphar.2023.1211663

Vega-Rivera, N.-M., Fernández-Guasti, A., Ramírez-Rodríguez, G., & Estrada-Camarena, E. (2014) Forced swim and chronic variable stress reduced hippocampal cell survival in OVX female rats. Behavioural Brain Research, 270, 248–55. 10.1016/j.bbr.2014.05.033

Walf, A. A., & Frye, C. A. (2007). The use of the elevated plus maze as an assay of anxiety-related behavior in rodents. Nature Protocols, 2(2), 322–328. 10.1038/nprot.2007.44

Yang, X., Fang, Y., Chen, H., Zhang, T., Yin, X., Man, J., Yang, L., & Lu, M. (2021). Global, regional and national burden of anxiety disorders from 1990 to 2019: Results from the Global Burden of Disease Study 2019. Epidemiology and Psychiatric Sciences, 30, e36. 10.1017/S2045796021000275

Zhu, X., Sun-Waterhouse, D., & Cui, C. (2021). A red pomegranate fruit extract-based formula ameliorates anxiety/depression-like behaviors *via* enhancing serotonin (5-HT) synthesis in C57BL/6 male mice. Food Science and Human Wellness, 10(3), 289–296. 10.1016/j.fshw.2021.02.020

